# Functionally-informed fine-mapping and polygenic localization of complex trait heritability

**DOI:** 10.1101/807792

**Authors:** Omer Weissbrod, Farhad Hormozdiari, Christian Benner, Ran Cui, Jacob Ulirsch, Steven Gazal, Armin P. Schoech, Bryce van de Geijn, Yakir Reshef, Carla Márquez-Luna, Luke O’Connor, Matti Pirinen, Hilary K. Finucane, Alkes L. Price

## Abstract

Fine-mapping aims to identify causal variants impacting complex traits. Several recent methods improve fine-mapping accuracy by prioritizing variants in enriched functional annotations. However, these methods can only use information at genome-wide significant loci (or a small number of functional annotations), severely limiting the benefit of functional data. We propose PolyFun, a computationally scalable framework to improve fine-mapping accuracy using genome-wide functional data for a broad set of coding, conserved, regulatory and LD-related annotations. PolyFun prioritizes variants in enriched functional annotations by specifying prior causal probabilities for fine-mapping methods such as SuSiE or FINEMAP, employing special procedures to ensure robustness to model misspecification and winner’s curse. In simulations with in-sample LD, PolyFun + SuSiE and PolyFun + FINEMAP were well-calibrated and identified >20% more variants with posterior causal probability >0.95 than their non-functionally informed counterparts (and >33% more fine-mapped variants than previous functionally-informed fine-mapping methods). In simulations with mismatched reference LD, PolyFun + SuSiE remained well-calibrated when reducing the maximum number of assumed causal SNPs per locus, which reduces absolute power but still produces large relative improvements. In analyses of 49 UK Biobank traits (average *N*=318K) with in-sample LD, PolyFun + SuSiE identified 3,025 fine-mapped variant-trait pairs with posterior causal probability >0.95, a >32% improvement vs. SuSiE; 223 variants were fine-mapped for multiple genetically uncorrelated traits, indicating pervasive pleiotropy. We used posterior mean per-SNP heritabilities from PolyFun + SuSiE to perform polygenic localization, constructing minimal sets of common SNPs causally explaining 50% of common SNP heritability; these sets ranged in size from 28 (hair color) to 3,400 (height) to 2 million (number of children). In conclusion, PolyFun prioritizes variants for functional follow-up and provides insights into complex trait architectures.

## Introduction

Genome-wide association studies of complex traits have been extremely successful in identifying loci harboring causal variants but less successful in fine-mapping the underlying causal variants, making the development of fine-mapping methods a key priority^1,2^. Fine-mapping methods aim to pinpoint causal variants by accounting for linkage disequilibrium (LD) between variants^3–12^, but have limited power in the presence of strong LD. One way to increase fine-mapping power is to prioritize variants in functional annotations that are enriched for complex trait heritability^7,8,10,13–17^. However, previous functionally-informed fine-mapping methods such as PAINTOR^18^, fastPAINTOR^19^, and CAVIARBF^20^ have computational limitations and can only use genome-wide significant loci to estimate functional enrichment (and the extension of fGWAS^21^ proposed in ref. ^10^ can only incorporate a small number of functional annotations), severely limiting the benefit of functional data.

We propose PolyFun, a computationally scalable framework for functionally-informed fine-mapping that makes full use of genome-wide data. PolyFun prioritizes variants in enriched functional annotations by defining prior causal probabilities for fine-mapping methods such as SuSiE^22^ or FINEMAP^23,24^. PolyFun estimates functional enrichment using a broad set of coding, conserved, regulatory, MAF and LD-related annotations from the baseline-LF model^25–27^, aggregating data from across the entire genome and hundreds of functional annotations, via a novel framework that incorporates stratified LD score regression^17^ and is robust to modeling misspecification and winner’s curse.

We show in simulations with in-sample LD that PolyFun is well-calibrated and is more powerful than previous fine-mapping methods, with a >20% power increase over non-functionally informed fine-mapping methods. In simulations with mismatched reference LD, PolyFun remains well-calibrated when reducing the maximum number of assumed causal SNPs per locus, which reduces absolute power but still produces large relative improvements over non-functionally informed fine-mapping methods. We apply PolyFun to 49 complex traits from the UK Biobank^28^ (average *N*=318K) with in-sample LD and identify 3,025 fine-mapped variant-trait pairs with posterior causal probability >0.95, spanning 2,225 unique variants. 223 of these variants were fine-mapped for multiple genetically uncorrelated traits, indicating pervasive pleiotropy. We further used posterior mean per-SNP heritabilities from PolyFun + SuSiE to perform polygenic localization, finding sets of common SNPs causally explaining 50% of common SNP heritability that range in size across many orders of magnitude, from dozens to hundreds of thousands of SNPs.

## Results

### Overview of methods

PolyFun prioritizes variants in enriched functional annotations by specifying prior causal probabilities in proportion to predicted per-SNP heritabilities and providing them as input to fine-mapping methods such as SuSiE^22^or FINEMAP^23,24^. For each target locus, PolyFun robustly specifies prior causal probabilities for all SNPs on the corresponding odd (resp. even) target chromosome by (1) estimating functional enrichments for a broad set of coding, conserved, regulatory and LD-related annotations from the baseline-LF 2.2.UKB model^26^ (187 annotations; Methods, Supplementary Table 1; see URLs) using an L2-regularized extension of S-LDSC^17^, restricted to even (resp. odd) chromosomes; (2) estimating per-SNP heritabilities for SNPs on odd (resp. even) chromosomes using the functional enrichment estimates from step 1; (3) partitioning all SNPs into 20 bins of similar estimated per-SNP heritabilities from step 2; (4) re-estimating per-SNP heritabilities for all SNPs on the target chromosome by applying S-LDSC to the 20 bins, restricted to odd (resp. even) chromosomes excluding the target chromosome; and (5) setting prior causal probabilities for SNPs on the target chromosome proportional to per-SNP heritabilities from step 4. The L2 regularization in step 1 improves the accuracy of per-SNP heritability estimation; the partitioning into odd and even chromosomes in steps 1-2 and the exclusion of the target chromosome in step 4 prevents winner’s curse; and the re-estimation of per-SNP heritabilities in step 4 ensures robustness to model misspecification. We evaluate the impact of each of these steps in the experiments below.

**Table 1:**
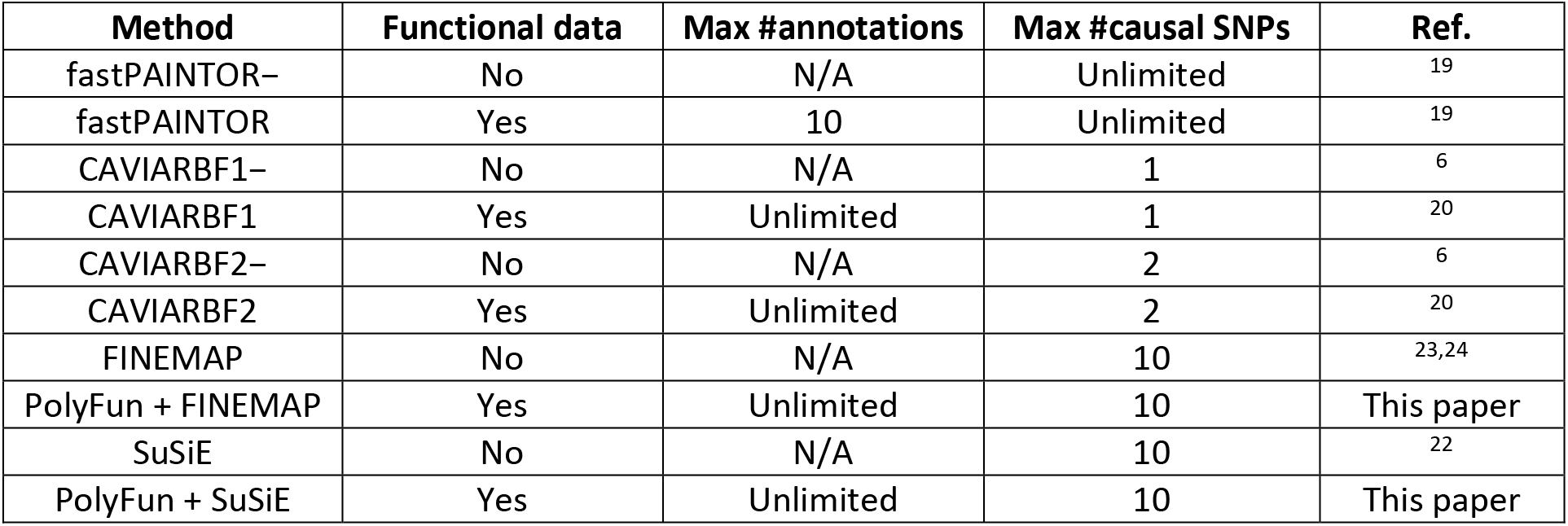
Summary of methods evaluated in main simulations. For each method we report whether it incorporates functional data, the maximum number of functional annotations that we specified under default simulation settings (for fastPAINTOR we selected the number of annotations that maximized power while maintaining correct calibration; Methods), the maximum number of causal SNPs modeled per locus (or the exact number for SuSiE and PolyFun + SuSiE), and the corresponding reference. For fastPAINTOR and CAVIARBF, − denotes the exclusion of func8onal data. For CAVIARBF, 1 or 2 denotes the maximum number of causal variants. PolyFun + FINEMAP uses a new version of FINEMAP that we introduce here that incorporates prior causal probabilities.

PolyFun specifies prior causal probabilities in proportion to per-SNP heritability estimates:

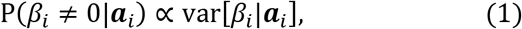

where *β*_*i*_ is the causal effect size of SNP *i* in standardized units (the number of standard deviations increase in phenotype per 1 standard deviation increase in genotype), ***α***_*i*_ is the vector of functional annotations of SNP *i*, and var[*β*_*i*_|***α***_*i*_] is the estimated per-SNP heritability of SNP *i* from step 4 (see above). Equation 1 is derived by applying the law of total variance to var[*β*_*i*_|***α***_*i*_], assuming that SNP effect sizes are sampled from a mixture distribution and that functional enrichment is primarily due to differences in polygenicity, motivated by our recent work^29^ (see Methods).

A key distinction between PolyFun and previous functionally-informed fine-mapping methods^10,18–20^ is the use of the entire genome and a large number of functional annotations to estimate prior causal probabilities. PolyFun achieves this by decoupling functional enrichment estimation and fine-mapping, which allows rapidly pooling data across millions of SNPs and >100 functional annotations from the baseline-LF model (see Methods). In contrast, previous functionally-informed fine-mapping methods can only aggregate information across a small number of loci (e.g. genome-wide significant loci)^18–20^ or a small number of functional annotations^10^ due to computational limitations. We exploited the computational scalability of PolyFun (together with SuSiE^22^) to fine-map up to 2,763 overlapping 3Mb loci spanning the entire genome (excluding loci with close to zero heritability; Methods), instead of only analyzing genome-wide significant loci. We subsequently used our fine-mapping results to perform polygenic localization, identifying minimal sets of common SNPs causally explaining a given proportion of common SNP heritability. Details of the PolyFun method are provided in the Methods section; we have released open-source software implementing PolyFun in conjunction with SuSiE22 and FINEMAP23 (see URLs). In all main simulations and analyses of real traits, we applied PolyFun using summary LD information estimated directly from the target samples (both for running S-LDSC and for running SuSiE or FINEMAP), as previously recommended for fine-mapping methods^12,30^. We also performed a comprehensive set of simulations to assess the impact of mismatched reference LD.

### Main simulations

We evaluated PolyFun via simulations using real genotypes from 337,491 unrelated UK Biobank samples of British ancestry^28^. We analyzed 10 3Mb loci (following previous recommendations on fine-mapping locus size^12^) on chromosome 1 with different SNP densities, each containing 1,468-27,784 imputed MAF≥0.001 SNPs (including short indels; Supplementary Table 2). We estimated prior causal probabilities using 18,212,157 genome-wide imputed MAF≥0.001 SNPs with INFO score≥0.6 (excluding three long-range LD regions; see Methods). We simulated traits with heritability equal to 25% and genome-wide proportion of causal SNPs equal to 0.5%, with the target locus in each simulation including 10 causal SNPs jointly explaining heritability equal to 0.05%. To define prior causal probabilities for the generative model, we generated a per-SNP heritability for every SNP based on its LD, MAF, and functional annotations, using the baseline-LF model^26^ with meta-analyzed functional enrichments from analyses of real data (Supplementary Table 3), and then defined causal probabilities in proportion to these per-SNP heritabilities (Equation 1; Methods). We sampled causal SNPs based on these causal probabilities, sampled causal effect sizes from the same normal distribution for each causal SNP (motivated by our recent work^29^), and generated summary statistics with sampling noise based on *N*=320K samples (corresponding to a 95% phenotyping rate for *N*=337K samples) using summary LD information from the same samples. Other parameter settings were explored in secondary analyses (see below). Further details of the simulation framework are provided in the Methods section.

**Table 2:** Pleiotropic fine-mapped SNPs for UK Biobank traits. We report SNPs fine-mapped (PIP>0.95) for ≥4 genetically uncorrelated traits (|*r_g_*|<0.2). For each SNP we report its name (SNP), position (hg19), MAF in the UK Biobank, closest gene(s) (using data from the GWAS catalog^80^), top annotation (Methods) and fine-mapped traits (and the number of fine-mapped traits). SNPs are ordered first by the number of fine-mapped traits and then by genomic position. HDL: HDL cholesterol; MC: monocyte count; MPV: mean platelet volume; HLSRC: high light scatter reticulocyte count; Cholesterol: total cholesterol; RBCDW: red blood cell distribution width; FEV1/FVC: ratio of forced expiratory volume to forced vital capacity; MCH: mean corpuscular hemoglobin; SBP: systolic blood pressure; DBP: diastolic blood pressure; FVC: forced vital capacity; Cardiovascular: cardiovascular-related disease; RBC: red blood cell count; LC: lymphocyte count; HbA1c: Hemoglobin A1c; WHR: waist-hip ratio (adjusted for BMI). Results for all 223 pleiotropic fine-mapped SNPs are reported in Supplementary Table 15.

| SNP | Position | MAF | Closest gene(s) | Annotation | Traits |
| --- | --- | --- | --- | --- | --- |
| rs13107325 | chr4:103188709 | 0.08 | SLC39A8 | non-synonymous | BMI, Balding, Cholesterol, DBP, FVC, Height, RBC, WHR (8) |
| rs1229984 | chr4:100239319 | 0.02 | ADH1B | non-synonymous | BMI, LDL, MCH, MPV, SBP, Vitamin D (6) |
| rs76895963 | chr12:4384844 | 0.02 | CCND2,CCND2-AS1 | Conserved | BMD, Height, RBC, SBP, Triglycerides (5) |
| rs140584594 | chr1:110232983 | 0.27 | GSTM1 | non-synonymous | HDL, Height, MC, MPV (4) |
| rs3811444 | chr1:248039451 | 0.33 | TRIM58 | non-synonymous | HLSRC, HbA1c, Platelet Count, RBC (4) |
| rs1260326 | chr2:27730940 | 0.39 | GCKR | non-synonymous | Cholesterol, Height, Platelet Count, RBCDW (4) |
| rs2270894 | chr3:9975386 | 0.2 | CRELD1,IL17RC | Conserved | BMD, FEV1/FVC, Height, Platelet Count (4) |
| rs11556924 | chr7:129663496 | 0.39 | ZC3HC1 | non-synonymous | Age Menarche, Cardiovascular, Height, Platelet Count (4) |
| rs3918226 | chr7:150690176 | 0.08 | NOS3 | Conserved | Eczema, Height, High Cholesterol, MPV (4) |
| rs150813342 | chr9:135864513 | 0.01 | GFI1B | Conserved | Eosinophil Count, HLSRC, MCH, Platelet Count (4) |
| rs964184 | chr11:116648917 | 0.13 | ZPR1 | DHS | Cholesterol, MPV, RBCDW, Vitamin D (4) |
| rs35979828 | chr12:54685880 | 0.07 | NFE2 | Conserved | Eosinophil Count, Platelet Count, RBC, RBCDW (4) |
| rs2277339 | chr12:57146069 | 0.1 | PRIM1 | non-synonymous | Height, LC, RBC, RBCDW (4) |
| rs72681869 | chr14:50655357 | 0.01 | SOS2 | non-synonymous | FVC, Hair Color, HbA1c, SBP (4) |
| rs61745086 | chr16:88782050 | 0.01 | PIEZO1,CTU2 | non-synonymous | HLSRC, HbA1c, Height, RBC (4) |
| rs34557412 | chr17:16852187 | 0.01 | TNFRSF13B | non-synonymous | HbA1c, MC, MPV, RBC (4) |
| rs77542162 | chr17:67081278 | 0.02 | ABCA6 | non-synonymous | HbA1c, Height, LDL, Platelet Count (4) |

**Table 3:** Summary of mismatched reference LD simulations. For each experiment (exp) we report: (GWAS) The sample size and population of the target sample (UK denotes British-ancestry individuals from UK Biobank; EUR denotes non-British European-ancestry individuals from UK Biobank, REL indicates that pairs of related individuals are included in the sample); (LD) the sample size and population of the LD reference panel (UK denotes British-ancestry individuals from UK Biobank; UK10K denotes individuals from the UK10K cohort; numbers in parentheses indicate how many individuals overlap the target sample, if any; “none” indicates that there is no LD reference panel); (Generative SNPs) The set of SNPs from which we sampled causal SNPs (UKB: the set of UK Biobank imputed SNPs with INFO score >0.6 and UKB MAF>0.1%; UK10K: the set of UK10K SNPs; INF: the set of UKB imputed SNPs with INFO score >0.9; COM: the set of UKB imputed SNPs with MAF >1% in British-ancestry individuals); (SNPs analyzed) the set of SNPs that was used for fine-mapping; and (max. *L*) The maximum number of causal SNPs per locus assumed by PolyFun + SuSiE that maintains FDR<0.05 at a PIP=0.95 threshold (selected from the options 1,2,3,10; - indicates that none of these options maintains FDR<0.05). Horizontal lines indicate the partitioning into types of experiments described in the main text. Numerical results are reported in Supplementary Table 7.

| exp | GWAS | LD | Generative SNPs | SNPs analyzed | max. $L$ |
| --- | --- | --- | --- | --- | --- |
| a | 44K UK | 44K UK (44K overlap) | UKB | UKB | 10 |
| b | 44K UK | 44K UK | UKB | UKB | 10 |
| c | 44K UK | 4K UK | UKB | UKB | 10 |
| d | 44K UK | 400 UK | UKB | UKB | 1 |
| e | 44K UK | none | UKB | UKB | 1 |
| f | 293K UK | 44K UK | UKB | UKB | 10 |
| g | 293K UK | 4K UK | UKB | UKB | 2 |
| h | 293K UK | 4K UK (4K overlap) | UKB | UKB | 2 |
| i | 44K EUR | 44K UK | UKB | UKB | 3 |
| j | 44K EUR | 4K UK | UKB | UKB | 2 |
| k | 44K EUR | 400 UK | UKB | UKB | 1 |
| l | 22K EUR+22K UK | 44K UK (22K overlap) | UKB | UKB | 3 |
| m | 44K UK-REL | 44K UK | UKB | UKB | 10 |
| n | 44K EUR-REL | 44K UK | UKB | UKB | 3 |
| o | 44K UK | 3.6K UK10K | UKB | UK10K∩UKB | - |
| p | 44K UK | 3.6K UK10K | UK10K∩UKB | UK10K∩UKB | 2 |
| q | 44K UK | 3.6K UK10K | UK10K∩UKB∩INF | UK10K∩UKB∩INF | 10 |
| r | 44K UK | 3.6K UK10K | UK10K∩UKB∩COM | UK10K∩UKB∩COM | 1 |
| s | 44K UK | 4K UK | UK10K∩UKB | UK10K∩UKB | 10 |

We evaluated 10 fine-mapping methods (Table 1): fastPAINTOR−, fastPAINTOR, CAVIARBF1−, CAVIARBF1, CAVIARBF2−, CAVIARBF2, FINEMAP, PolyFun + FINEMAP, SuSiE, and PolyFun + SuSiE. (We did not evaluate methods that do not incorporate functional annotations or prior causal probabilities, such as CAVIAR^5^.) For each method, we used summary LD information estimated directly from the target samples, as in our analyses of real traits. For fastPAINTOR−, fastPAINTOR, SuSiE, and PolyFun + SuSiE, we specified a per-locus causal effect size variance (var[*β*_*i*_|*β*_*i*_ ≠ 0]) using a causal effect size variance estimator that we implemented, which yielded improved results over the default estimator implemented in these methods (see Methods). We ran fastPAINTOR with 10 annotations that we selected to maximize power while maintaining correct calibration (Methods). We used default settings for all other parameters, except as otherwise indicated. We applied fastPAINTOR, CAVIARBF1, and CAVIARBF2 to one locus at a time due to computational limitations (see below). Results for FINEMAP, PolyFun + FINEMAP, SuSiE, and PolyFun + SuSiE were averaged across 1,000 simulations, and results for other methods were averaged across 100 simulations due to computational limitations.

We assessed calibration via the proportion of false positives among SNPs with posterior causal probability (posterior inclusion probability; PIP) above a given threshold (e.g. PIP>0.95 or PIP>0.5), aggregating the results across all simulations; we refer to this quantity as the false discovery rate (FDR) although we do not use frequentist false discovery rate methods^31,32^. For each PIP threshold, we conservatively estimated the FDR as one minus the PIP threshold, which can be motivated under a Bayesian interpretation of FDR (Methods, Figure 1a-b, Supplementary Tables 4). (We alternatively explored a data-driven estimator defined as one minus the average PIP among all SNPs with PIP greater than the threshold and found that it is anti-conservative and hence not recommended (Methods, Figure 1a-b, Supplementary Tables 4), possibly because computing exact PIPs is computationally intractable or because of imperfectly-estimated priors, demonstrating the challenges of exact calibration in fine-mapping). No method except CAVIARBF2− and CAVIARBF2 had significantly inflated false discovery rates, although fastPAINTOR and CAVIARBF1 had suggestive evidence of inflated false discovery rates (due to high computational costs, we did not perform enough simulations to determine if this miscalibration was statistically significant). CAVIARBF2− and CAVIARBF2 had severely inflated false discovery rates at multiple PIP cutoffs (Supplementary Table 4), contrary to expectations as modeling multiple causal variants is advantageous in most settings^3^. Similarly, no method had a significantly inflated false discovery rate at the PIP>0.95 threshold when using exact false-discovery thresholds, although all methods had inflated false discovery rates at the PIP>0.5 threshold with exact false-discovery thresholds (significant for methods for which we ran 1,000 simulations), demonstrating the challenges of exact calibration in fine-mapping (Methods, Figure 1, Table 4).

**Figure 1:**
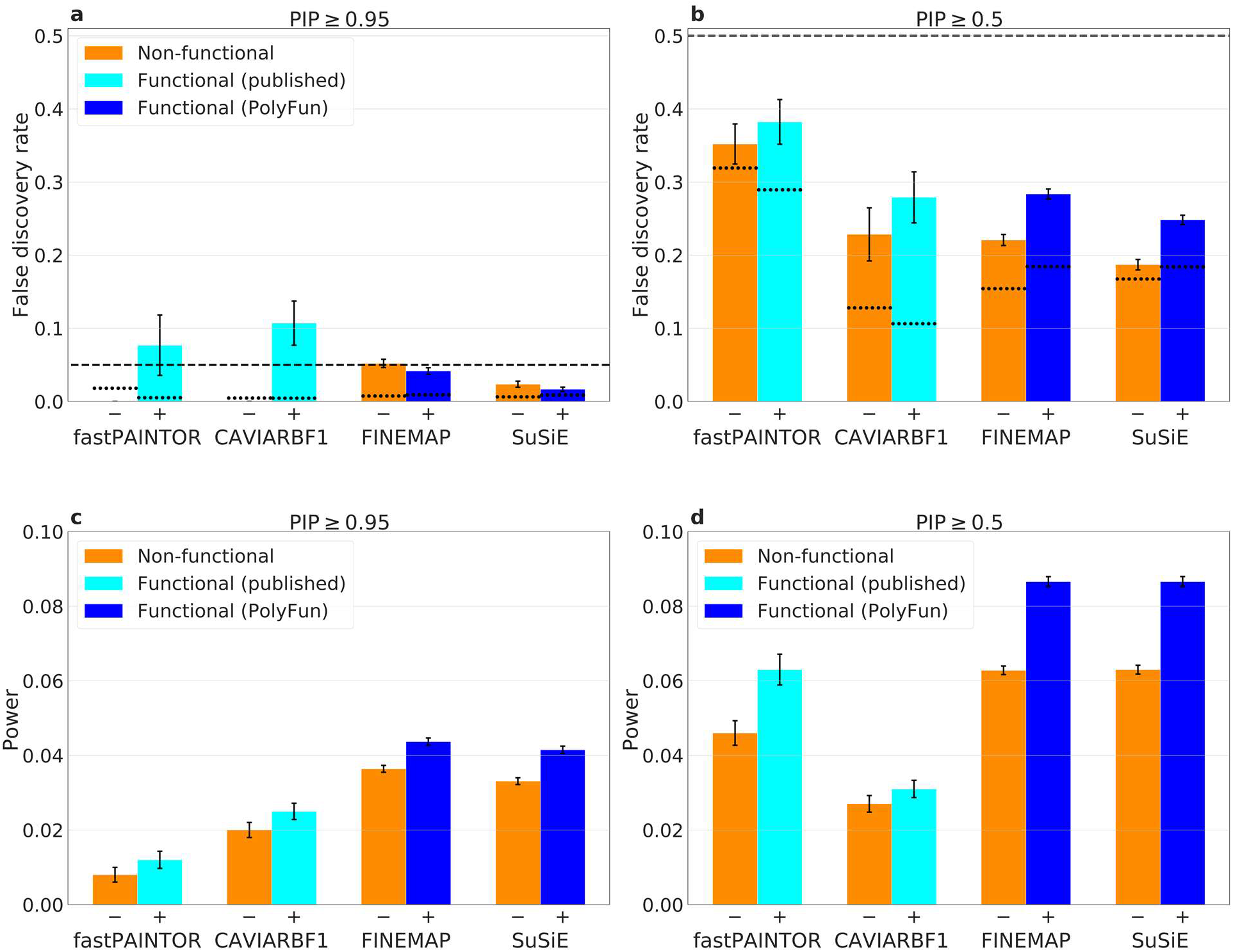
Calibration and power of fine-mapping methods in main simulations. **(a-b) FDR at PIP=0.95 (a) and PIP=0.5 (b)**. Upper dashed horizontal lines denote conservative FDR estimates. Lower dotted horizontal lines denote anti-conservative FDR estimates, which are not recommended (Methods). (c-d) Power at PIP=0.95 (c) and PIP=0.5 (d). The first bar of each method uses non-functionally informed fine-mapping (denoted −), and the second uses func8onally informed fine-mapping (denoted +). Error bars denote standard errors. Numerical results, including results for CAVIARBF2− and CAVIARBF2, are reported in Supplementary Table 4.

We assessed power via the proportion of true causal SNPs with PIP above a given threshold, aggregating the results across all simulations (Figure 1c-d, Supplementary Table 4). PolyFun + FINEMAP was the most powerful method, identifying >5% more PIP>0.95 causal SNPs than PolyFun + SuSiE and >20% more PIP>0.95 causal SNPs than FINEMAP (Supplementary Table 4). PolyFun + SuSiE was the second most powerful method, identifying >25% more PIP>0.95 causal SNPs than SuSiE. PolyFun + FINEMAP and PolyFun + SuSiE were equally powerful at PIP>0.5, identifying >37% more PIP>0.5 causal SNPs than all other methods. These results demonstrate the benefits of using functional annotations for SNP prioritization. We note that power to identify PIP>0.95 or PIP>0.5 SNPs is generally expected to be low (on the order of 10% or lower), as fine-mapping is a statistically hard problem due to pervasive LD^3^. However, power was substantially higher at lower PIP thresholds (Supplementary Table 4).

We evaluated the computational cost of each method. SuSiE and PolyFun + SuSiE were much faster than the other methods, fine-mapping a 3Mb locus in 5 minutes on average (excluding fixed preprocessing time; see below), compared with 227 minutes for fastPAINTOR−, 235 minutes for fastPAINTOR, 20 minutes for CAVIARBF1−, 34 minutes for CAVIARBF1, 70 minutes for CAVIARBF2−, 84 minutes for CAVIARBF2, 14 minutes for FINEMAP, and 17 minutes for PolyFun + FINEMAP (Figure 2a, Supplementary Table 4). CAVIARBF methods allowing >2 causal SNPs per locus were not evaluated because they typically required >48 hours for a single analysis. We note that the computational costs for fastPAINTOR, CAVIARBF1, and CAVIARBF2 that we report here for 3Mb loci are larger than those previously reported for smaller loci (typically ≤100kb)^18–20^. PolyFun also requires fixed preprocessing time (steps 1-4; see Overview of methods) of 630 minutes on average (equivalent to 0.2 minutes per locus in a genome-wide analysis); when restricting analyses to subsets of loci, PolyFun + SuSiE was still faster than all other functionally-informed methods when analyzing >23 loci (Figure 2b).

**Figure 2:**
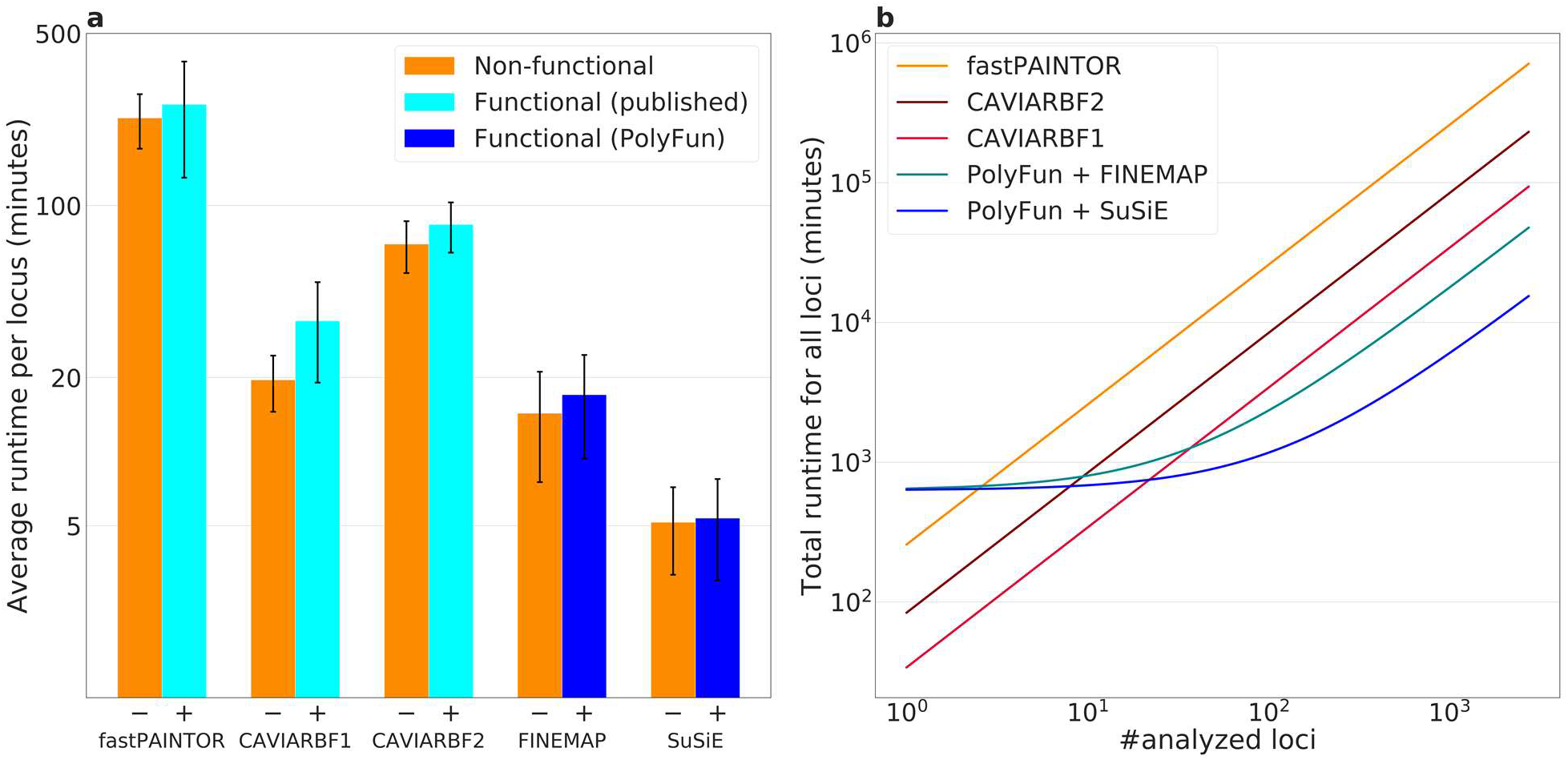
Computational cost of fine-mapping methods. (**a**) The average runtime required to fine-map a 3Mb locus in a genome-wide analysis (log scale). The first bar of each method uses non-functionally informed fine-mapping (denoted −), and the second uses functionally informed fine-mapping (denoted +). (**b**) The total runtime required to fine-map different numbers of loci, for functionally informed fine-mapping methods only (log scale). The runtimes of PolyFun + SuSiE and PolyFun + FINEMAP are sub-linear because they include the fixed preprocessing cost of computing prior causal probabilities (630 minutes). Numerical results, including panel (b) results for non-functionally informed methods, are reported in Supplementary Table 4.

To assess the robustness of PolyFun to model misspecification of functional architectures (i.e. generative models of causal effect sizes as a function of functional annotations), we also simulated data using two functional architectures that are different from the additive functional architecture assumed by S-LDSC with the baseline-LF model^26^: (i) a multiplicative functional architecture in which per-SNP heritabilities are proportional to a product of terms for each annotation^21,33^, and (ii) a sub-additive functional architecture in which per-SNP heritabilities are proportional to a maximum of terms for each annotation (see Methods). In both cases, PolyFun + SuSiE and PolyFun + FINEMAP remained well-calibrated and attained a >24% increase in power over the respective non-functionally informed methods (Supplementary Table 4). On the other hand, alternative functionally-informed extensions of SuSiE and FINEMAP that specify prior causal probabilities (Equation 1) in proportion to standard S-LDSC per-SNP heritability estimates or to L2-regularized S-LDSC per-SNP heritability estimates (the output of step 1 of PolyFun) suffered inflated false discovery rates or reduced power (Supplementary Table 4), demonstrating the importance of robustly specifying prior causal probabilities.

We performed 5 additional experiments to assess the individual impact of each of steps 1-5 of the PolyFun method on fine-mapping performance. The experiments are summarized in Supplementary Table 5, and results are reported in Supplementary Table 6. First, we perturbed step 1 (estimating functional enrichment) by randomly flipping the sign of either 25%, 50%, or 75% of the annotation coefficient estimates 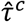. We determined that this led to significantly increased FDR (Supplementary Table 6a). Second, we perturbed step 2 (estimating per-SNP heritabilities on odd/even chromosomes) by first estimating annotation coefficients using both and even chromosomes in step 1, and then estimating per-SNP heritabilities for each SNP using these annotation coefficients. We determined that this led to significantly increased FDR and significantly reduced power (Supplementary Table 6b). Third, we perturbed step 3 (partitioning all SNPs into 20 bins of similar per-SNP heritability) by varying the number of bins to be either 5, 10, or 30. We determined that this led to no significant change in FDR and that using only 5 bins led to significantly decreased power but that using 10 or 30 bins led to no significant change in power (Supplementary Table 6c). Fourth, we perturbed step 4 (re-estimating per-SNP heritabilities within each bin excluding the target chromosome) by not excluding the target chromosome. We determined that this did not lead to statistically significant differences in the results under our default simulation settings. However, when repeating this experiment in simulations with smaller sample size (*N*=10K), this perturbation led to significantly increased FDR (Supplementary Table 6d). Fifth, we perturbed step 5 (specifying prior causal probabilities in proportion of the re-estimated per-SNP heritabilities) by randomly permuting either 25%, 50%, or 75% of the re-estimated per-SNP heritabilities before computing prior causal probabilities. We determined that this led to significantly increased FDR and significantly reduced power (Supplementary Table 6e). We conclude that each of steps 1-5 of the PolyFun method is crucial for controlling FDR and maximizing power.

We performed 9 secondary analyses. First, we evaluated 95% credible sets, defined as sets with probability >0.95 of including ≥1 causal SNP(s)^22^ (this is different from an earlier definition of 95% credible set as the smallest set of SNPs accounting for 95% of the posterior probability^4,5,18^); we note that 95% credible sets generally contain many non-causal SNPs. The union of PolyFun + SuSiE credible sets included 25 SNPs on average (27% fewer than SuSiE) and spanned 22% of the true causal SNPs (1% more than SuSiE) (Supplementary Table 4). The union of PolyFun + FINEMAP 95% credible sets included 27 SNPs on average (27% fewer than FINEMAP) and spanned 27% of the true causal SNPs (5% more than FINEMAP) (Supplementary Table 4). Second, we evaluated the methods under different simulated sample sizes. The relative power advantage of PolyFun + SuSiE over SuSiE (resp. PolyFun + FINEMAP over FINEMAP) at a PIP>0.95 threshold was equal to 22% (resp. 14%) at *N*=1 million, vs. 25% (resp. 20%) at *N*=320K (corresponding to the typical number of phenotyped individuals in the full UK Biobank), with a roughly linear increase in absolute power (Supplementary Table 4). These results indicate that functionally informed fine-mapping remains valuable at substantially larger sample sizes. Third, we verified that the improvement of PolyFun remained qualitatively similar for different values of the number of causal SNPs per target locus (Supplementary Table 4) or the heritability causally explained by the target locus (Supplementary Table 4). Fourth, we verified that the improvement of PolyFun remained qualitatively similar for different values of genome-wide polygenicity (Supplementary Table 4) and genome-wide heritability (Supplementary Table 4). Fifth, we verified that the improvement of PolyFun remained qualitatively similar when changing the number of causal SNPs per locus assumed by PolyFun + SuSiE (or the maximum number of causal SNPs per locus assumed by PolyFun + FINEMAP) from 10 to 1-5, while the absolute power of all the methods decreased (Supplementary Table 4). Sixth, for all methods not based on CAVIARBF, we compared the performance of our per-locus causal effect size variance estimator to the default estimators (CAVIARBF-based methods perform fine-mapping using several prespecified values and then average the results^20^). We determined that the false discovery rate of fastPAINTOR-, fastPAINTOR, SuSiE and PolyFun + SuSiE improved when using our estimator (Supplementary Table 4). On the other hand, for FINEMAP and PolyFun + FINEMAP, causal discovery rate and power remained similar when using our estimator, but the sizes of 95% credible set sizes increased substantially (Supplementary Table 4), and thus we used the default FINEMAP estimator in all primary analyses. Seventh, we verified that fastPAINTOR performance was approximately optimized when incorporating 10 functional annotations (selected to maximize power while maintaining correct calibration, see Methods) (Supplementary Table 4). Eighth, we determined that the false discovery rate of PolyFun increased when using unregularized S-LDSC in step 1 of PolyFun, demonstrating the importance of regularization for functionally-informed fine-mapping (Supplementary Table 4). Finally, we evaluated fine-mapping performance when specifying the true prior causal probabilities that we used to generate the data, a “cheating” method, and determined that this substantially reduced the false discovery rate and increased the power of PolyFun + SuSiE and PolyFun + FINEMAP (Supplementary Table 4), confirming that more accurate prior causal probabilities lead to more powerful fine-mapping.

We conclude from these experiments that PolyFun + FINEMAP and PolyFun + SuSiE outperformed all other methods, with a 3.4x faster runtime for PolyFun + SuSiE; fastPAINTOR−, fastPAINTOR, CAVIARBF1− and CAVIARBF1 had lower power and high computational costs; CAVIARBF2− and CAVIARBF2 had extremely inflated false discovery rates; and CAVIARBF methods allowing >2 causal SNPs per locus had prohibitive computational costs. (We note that the power of fastPAINTOR, CAVIARBF1, and CAVIARBF2 could potentially be improved by jointly fine-mapping multiple genome-wide significant loci, but the computational costs would be prohibitive when there are many such loci.) Thus, we restricted our analyses in the remainder of this manuscript to SuSiE and PolyFun + SuSiE.

### Simulations with mismatched reference LD

Our main simulations used in-sample LD computed directly from the target samples. Although we have publicly released summary LD information for British-ancestry UK Biobank samples as part of this study (see URLs), there are many settings in which researchers conducting fine-mapping cannot obtain in-sample LD, and instead use LD information from an external LD reference panel^34^. We performed extensive simulations to assess how fine-mapping performance is impacted by LD mismatch between the target sample and the LD reference panel. We specifically considered (1) non-overlapping target and reference samples; (2) sample sizes of the target sample and reference panel; (3) differences in ancestry; (4) presence of related individuals in the target sample; and (5) SNPs available for analysis in the target sample and reference panel.

PolyFun + SuSiE and other fine-mapping methods are unaffected by mismatched reference LD when assuming a single causal SNP per locus, because they do not make use of summary LD information in this setting (Supplementary Note), but assuming a single causal SNPs per locus can decrease power when there are multiple causal SNPs. More generally, increasing the assumed number of causal SNPs increases the power of PolyFun + SuSiE when there are multiple causal SNPs (Supplementary Table 4). However, increasing the assumed number of causal SNPs can increase the false discovery rate, exacerbating the effects of LD mismatch. Hence, we quantified how mismatched reference LD impacts fine-mapping performance via the maximum number of assumed causal SNPs per locus that maintains FDR<0.05 at a PIP=0.95 threshold.

Our mismatched reference LD simulations differed from our main simulations in the following ways: we generated summary statistics using up to *N*=44K unrelated (or related) European-ancestry (British or non-British) UK Biobank target samples in most experiments, compared with *N*=320K in our main simulations, because the UK Biobank includes only 44K unrelated UK Biobank individuals of non-British European ancestry (we used *N*=293K unrelated British-ancestry UK Biobank target samples in a subset of experiments to more closely match our main simulations); we computed summary LD information using either *N*=400, *N*=4,000, or *N*=44K unrelated British-ancestry UK Biobank reference samples (either non-overlapping or overlapping with the target samples), or using *N*=3,567 reference samples from the UK10K cohort^35^; we generated summary statistics using raw genotypes rather than summary LD information (as required when the target sample and the LD reference panel are not the same); we simulated 3 causal SNPs per locus that jointly explain 0.5% of trait variance, compared with 10 causal SNPs that jointly explain 0.05% of trait variance in our main simulations, to obtain sufficient power despite having a smaller sample size; and in some experiments we used a subset of SNPs for generating causal SNPs or for fine-mapping analysis.

We performed 19 experiments, averaging the results of each experiment across 1,000 simulations (Table 3). For each experiment we report the maximal number of causal SNPs per locus assumed by PolyFun + SuSiE (denoted as *L*), that maintains FDR<0.05 at PIP=0.95. The target sample and the LD reference panel did not overlap unless stated otherwise. Our results are summarized in Table 3 and described in detail below.

We performed 2 experiments to evaluate the impact of non-overlapping target and reference samples of the same ancestry and sample size. First, we performed simulations analogous to our main simulations (Figure 1) but at lower sample size: we used an LD reference panel of *N*=44K unrelated British-ancestry UK-Biobank individuals and a target sample of the same individuals, which maintained FDR<0.05 for *L*≤10 (Supplementary Table 7a). Second, we used a target sample of *N*=44K unrelated British-ancestry UK Biobank individuals that did not overlap the LD reference panel, which maintained FDR<0.05 for *L*≤10 with comparable power (Supplementary Table 7b). We conclude that PolyFun + SuSiE is not impacted by non-overlapping target and reference samples of the same ancestry and sample size.

We performed 6 experiments to evaluate the impact of sample size differences between ancestry-matched target and reference samples. First, we decreased the LD reference panel size to *N*=4K, which maintained FDR<0.05 for *L*≤10 (Supplementary Table 7c). Second, we decreased the LD reference panel size to *N*=400, which maintained FDR<0.05 for *L*=1 only (Supplementary Table 7d). Third, we decreased the LD reference panel size to *N*=0 (using an LD matrix that is equal to the identity matrix), which maintained FDR<0.05 for *L*=1 only (Supplementary Table 7e). Fourth, we used a target sample of *N*=293K unrelated British-ancestry UK Biobank individuals, which maintained FDR<0.05 for *L*≤10 (Supplementary Table 7f). Fifth, we used the same *N*=293K target sample with a *N*=4K LD reference panel, which maintained FDR<0.05 for *L*≤2 only (Supplementary Table 7g). Sixth, we used the same *N*=293K target sample with a *N*=4K LD reference panel that overlapped the target sample, which maintained FDR<0.05 for *L*≤2 only (Supplementary Table 7h). We conclude that PolyFun + SuSiE with *L*=10 is well-calibrated if the LD reference panel is ancestry-matched to the target sample and spans ≥10% of the target sample size.

We performed 4 experiments to evaluate the impact of ancestry differences between the target and reference samples. First, we used an LD reference panel of *N*=44K unrelated British-ancestry UK Biobank individuals and a target sample of *N*=44K unrelated Non-British European ancestry UK Biobank individuals, which maintained FDR<0.05 for *L*≤3 (Supplementary Table 7i). Second, we decreased the LD reference sample size to *N*=4K, which maintained FDR<0.05 for *L*≤2 (Supplementary Table 7j). Third, we decreased the LD reference panel size to *N*=400, which maintained FDR<0.05 for *L*=1 only (Supplementary Table 7k). Fourth, we used a target sample of *N*=44K unrelated European-ancestry UK Biobank individuals, 22K of which overlapped the LD reference panel and the other 22K were of non-British ancestry, which maintained FDR<0.05 for *L*≤3 (Supplementary Table 7l). We conclude that when the target and reference sample are of different ancestries, PolyFun + SuSiE may not be well-calibrated under *L*>1.

We performed 2 experiments to evaluate the impact of the presence of related individuals in the target sample. First, we used a target sample of *N*=44K British-ancestry UK Biobank individuals without restricting to unrelated individuals, which maintained FDR<0.05 for *L*≤10 (Supplementary Table 7m). Second, we used a target sample of *N*=44K Non-British European-ancestry UK Biobank individuals without restricting to unrelated individuals, which maintained FDR<0.05 for *L*≤3 only (Supplementary Table 7n), comparable to the analogous experiment that restricted to unrelated individuals (Supplementary Table 7i). We conclude that PolyFun is robust to the presence of related individuals in the target sample, attaining essentially the same results as in experiments without related individuals; we note that these results apply to the typical levels of relatedness observed in UK Biobank and may not be applicable to cohorts with high levels of relatedness.

We performed 5 experiments to evaluate the impact of SNPs available for analysis in the target and reference samples. First, we used an LD reference panel of *N*=3,567 individuals from the UK10K cohort and restricted the set of SNPs considered in fine-mapping to the intersection of SNPs found in both UK10K and the UK Biobank, which yielded FDR>0.05 regardless of L because not all causal SNPs were considered in the fine-mapping analysis (Supplementary Table 7o). Second, we also restricted the set of SNPs of the generative model to the intersection of SNPs found in UK10K and the UK Biobank, which maintained FDR<0.05 for *L*≤2 only (Supplementary Table 7p). Third, we restricted the generative model and the fine-mapping analysis to use only SNPs found at the intersection the UK Biobank and UK10K and having INFO score>0.9, which maintained FDR<0.05 for *L*≤10 (Supplementary Table 7q). Fourth, we restricted the generative model and the fine-mapping analysis to use only SNPs found at the intersection the UK Biobank and UK10K and having MAF >1%, which maintained FDR<0.05 for *L*≤1 only (Supplementary Table 7r). Fifth, to verify that PolyFun + SuSiE could remain well-calibrated when restricting to UK10K SNPs, we used an LD reference panel of *N*=4K unrelated British-ancestry UK Biobank individuals while using only SNPs found in both UK10K and the UK Biobank for both the generative model and for fine-mapping analysis, which maintained FDR<0.05 for *L*≤10 (Supplementary Table 7s). We conclude that (i) fine-mapping analysis can yield false positive results if causal SNPs are not included in the LD reference panel; and (ii) there exist discrepancies in the LD patterns of INFO<0.9 SNPs between the UK10K cohort and British-ancestry individuals from the UK Biobank. These differences may stem either from subtle population differences between these two cohorts^36^ or from imputation errors (although a recent study of whole exome sequencing data from the UK Biobank indicates that the set of SNPs used in this manuscript, having INFO>0.6 and MAF>0.1%, is well imputed^37^).

Based on these experiments we provide the following fine-mapping best-practice recommendations: (1) PolyFun + SuSiE should ideally use in-sample LD from the GWAS target sample, with *L*=10; we have facilitated this option for UK Biobank researchers by publicly releasing summary LD information for British-ancestry UK Biobank samples as part of this study (see URLs); (2) PolyFun + SuSiE can alternatively use a non-overlapping LD reference panel from the target population that spans ≥10% of the target sample size, with *L*=10; (3) When there is no reference panel from the exact same population as the target sample that spans ≥10% of the target sample size, PolyFun + SuSiE can be used without an LD reference panel by specifying *L*=1 (Supplementary Note). We caution that even subtle population differences may lead to false positive results with *L*>1, as demonstrated in our analyses using UK10K as an LD reference panel.

Importantly, PolyFun + SuSiE holds a significant power advantage over other methods when specifying *L*=1, yielding >10% power gains while maintaining FDR<0.05 (compared with >20% power gains when using in-sample LD and specifying *L*=10, as reported in the main simulations); (4) PolyFun + SuSiE can be used in the presence of related individuals in the target sample, attaining essentially the same results with or without omitting related individuals from the sample; we note that these results apply to the typical levels of relatedness observed in UK Biobank and may not be applicable to cohorts with high levels of relatedness; and (5) PolyFun + SuSiE should include as many well-imputed SNPs from the target locus as possible to minimize the risk of omitting causal SNPs from the fine-mapping analysis.

### Functionally informed fine-mapping of 49 complex traits in the UK Biobank

We applied PolyFun + SuSiE to fine-map 49 traits in the UK Biobank, including 31 traits analyzed in refs. ^38,39^, 9 blood cell traits analyzed in ref. ^12^, and 7 recently released metabolic traits (average *N*=318K; Supplementary Table 8). For each trait we fine-mapped up to 2,763 overlapping 3Mb loci spanning M=18,212,157 imputed MAF≥0.001 SNPs with INFO score≥0.6 (including short indels; excluding three long-range LD regions and loci with close to zero heritability; Methods). We assigned to each SNP its PIP computed using the locus in which it was most central. We have publicly released the PIPs and the prior and posterior means and variances of the causal effect sizes for all SNPs and traits analyzed (see URLs).

PolyFun + SuSiE identified 3,025 PIP>0.95 fine-mapped SNP-trait pairs, a >32% improvement vs. SuSiE; 9,684 PIP>0.5 SNP-trait pairs, a >59% improvement vs. SuSiE; and 225,153 PIP>0.05 SNP-trait pairs, a >84% improvement vs. SuSiE (Supplementary Table 9). The number of PIP>0.95 SNPs per trait ranged from 0 (number of children) to 407 (height) (Figure 3a, Supplementary Table 9). The 3,025 PIP>0.95 SNP-trait pairs spanned 2,225 unique SNPs, including 532 low-frequency SNPs (0.005<MAF<0.05) and 185 rare SNPs (0.001<MAF<0.005) (Supplementary Table 10). Only 39% of the 2,225 PIP>0.95 SNPs were also lead GWAS SNPs (defined as MAF>0.001 SNPs with P<5×10^−8^ and no MAF>0.001 SNP with a smaller p-value within 1Mb) (Supplementary Table 10), demonstrating the importance of using fine-mapped SNPs rather than lead GWAS SNPs for downstream analysis. 31% of the PIP>0.95 SNPs resided in coding regions and 22% were non-synonymous (broadly consistent with previous fine-mapping studies^8,12^) (Supplementary Table 10). When restricting the analysis to 16 genetically uncorrelated traits (|*r_g_*|<0.2; Methods and Supplementary Tables 11-12) we identified 1,626 PIP>0.95 SNP-trait pairs spanning 1,496 unique SNPs, with a median distance of 9kb between a PIP>0.95 SNP and the nearest lead GWAS SNP for the same trait (Supplementary Table 10). The 16 genetically uncorrelated traits included 5,314 genome-wide significant locus-trait pairs (defined by 1Mb windows around lead GWAS SNPs) harboring 0.28 PIP>0.95 SNPs per locus on average (Supplementary Table 13); 9% of the PIP>0.95 SNP-trait pairs did not lie within genome-wide significant loci. 1,080 of the 5,306 locus-trait pairs (20%) harbored ≥1 PIP>0.95 SNP(s), harboring 1.37 PIP>0.95 SNPs on average (Supplementary Table 13).

**Figure 3:**
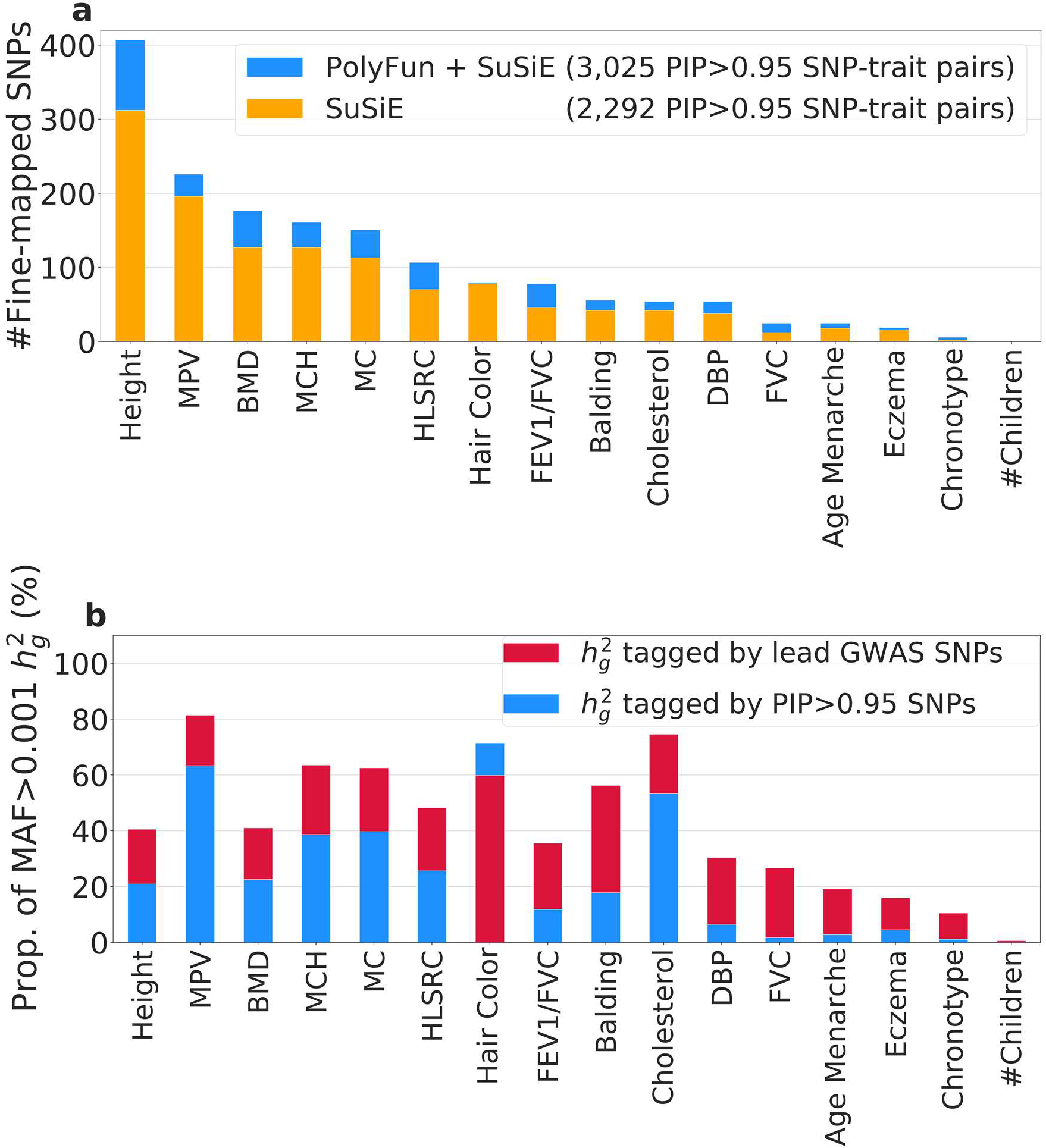
Summary of fine-mapping results for UK Biobank traits. (**a**) the number of SNPs with PIP>0.95 identified by SuSiE (orange bars) and PolyFun + SuSiE (blue bars) across 16 genetically uncorrelated traits in the UK Biobank. Traits are ordered by PolyFun + SuSiE results. The numbers in the legend refer to the sum of all 49 traits analyzed. (**b**) The proportion of MAF>0.001 SNP-heritability 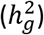 tagged by lead GWAS SNPs (crimson bars) and by PolyFun + SuSiE PIP>0.95 SNPs (blue bars). Traits are ordered as in panel (a). For hair color, the 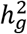 tagged by PIP>0.95 SNPs is greater than 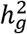 tagged by lead GWAS SNPs. MPV: Mean platelet volume; BMD: bone mineral density; MCH: mean corpuscular hemoglobin; MC: monocyte count; HLSRC: high light scatter reticulocyte count; FEV1/FVC: ratio of forced expiratory volume to forced vital capacity; DBP: diastolic blood pressure; FVC: forced vital capacity. Numerical results are reported in Supplementary Tables 9,14.

We estimated the SNP-heritability 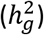 tagged by PIP>0.95 fine-mapped SNPs (which is likely to be close to the heritability causally explained by these SNPs, if most of the tagged SNP-heritability originates from PIP>0.95 SNPs). The 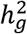 tagged by PIP>0.95 SNPs captured a large proportion of the 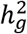 tagged by lead GWAS SNPs (median proportion=42%; Figure 3b, Methods, Supplementary Table 14). This proportion was substantially larger than the proportion of GWAS loci harboring PIP>0.95 SNPs (20%; see above), as fine-mapping power is higher at loci with larger causal effects (Supplementary Table 4). However, fine-mapped SNPs tagged a smaller proportion of total MAF>0.001 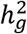 (median proportion=19%; Figure 3b, Methods, Supplementary Table 14), indicating that substantially larger sample sizes are required to comprehensively fine-map all heritable SNP effects.

Among the 2,225 unique PIP>0.95 SNPs fine-mapped for at least one trait, 223 SNPs were fine-mapped for multiple genetically uncorrelated traits (selecting a different subset of genetically uncorrelated traits for each SNP; Methods), including 55 SNPs fine-mapped for ≥3 genetically uncorrelated traits, indicating pervasive pleiotropy (Figure 4, Supplementary Table 15). 118 pleiotropic SNPs resided in coding regions and 93 were non-synonymous (Supplementary Table 15). The 17 SNPs fine-mapped for at least 4 traits are reported in Table 2. Top pleiotropic SNPs included (1) rs13107325, a non-synonymous SNP in the gene SLC39A8, which was fine-mapped for balding, BMI, diastolic blood pressure, forced vital capacity, height, red blood cell count, total cholesterol, and waist hip ratio (adjusted for BMI); (2) rs1229984, a non-synonymous SNP in the gene ADH1B, which was fine-mapped for BMI, LDL, mean corpuscular hemoglobin, mean platelet volume, systolic blood pressure, total cholesterol, and Vitamin D; and (3) rs76895963, a conserved intronic SNP in a promoter of the gene CCND2, which was fine-mapped for bone mineral density, height, red blood cell count, systolic blood pressure and triglycerides. The gene SLC39A8 is a zinc transporter and is associated with congenital disorder of glycosylation^40,41^; the gene ADH1B is an alcohol dehydrogenase gene and is associated with alcohol dependence^42^; and the gene CCND2 participates in cell cycle regulation and is associated with delayed psychomotor development^43^. We note that previous studies have reported that genetically uncorrelated traits often share association signals at the same loci^44^, but did not fine-map those signals to individual SNPs as performed here.

**Figure 4:**
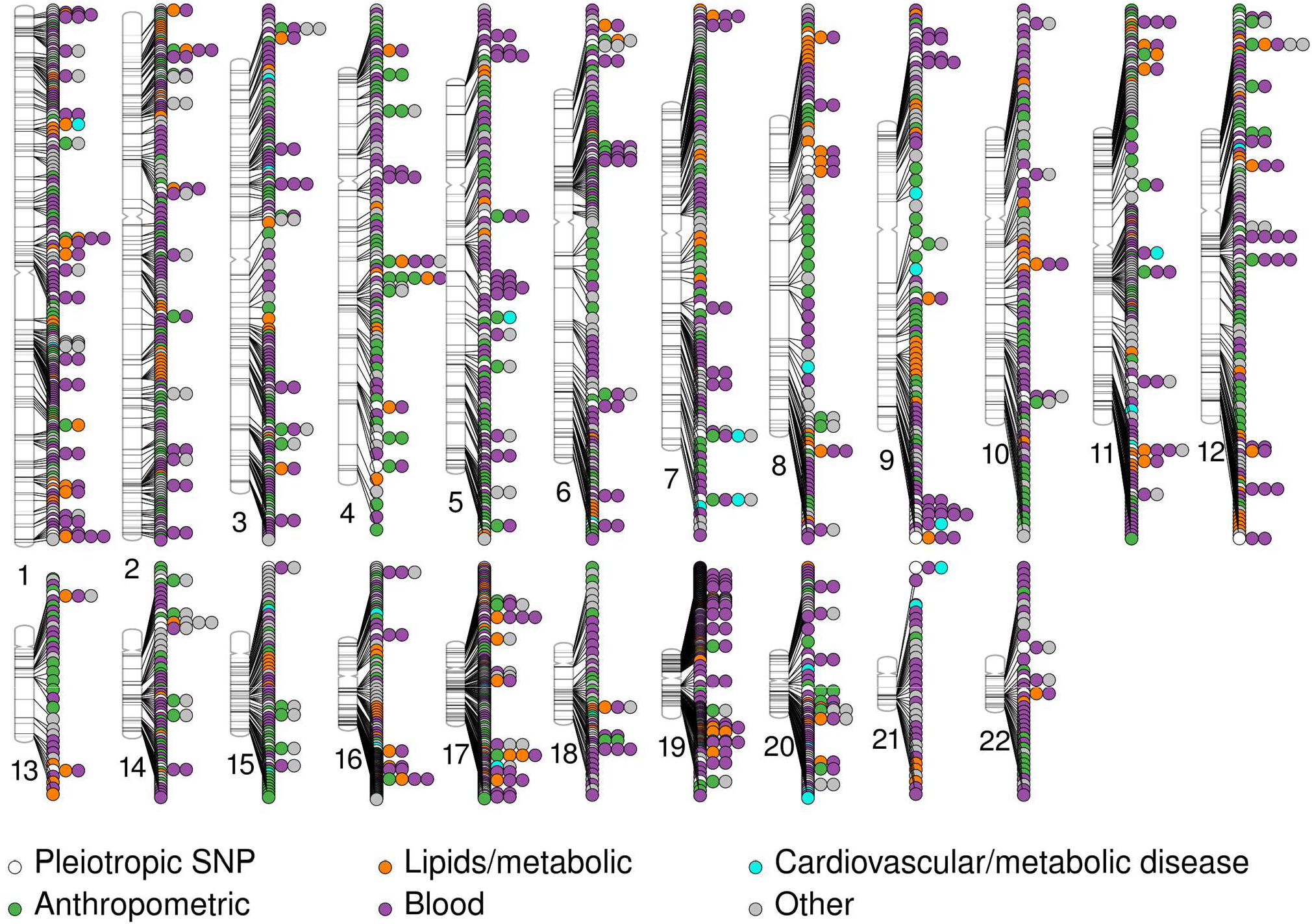
Visualization of fine-mapping results for UK Biobank traits. We display an ideogram of all 2,225 PIP>0.95 fine-mapped SNPs identified by PolyFun + SuSiE across 49 UK Biobank traits. Traits are color-coded into groups (see legend and Supplementary Table 8). White circles indicate SNPs that are pleiotropic for ≥2 genetically uncorrelated traits, with circles to the right of a white circle denoting the genetically uncorrelated traits (max of 5 colored circles due to space limitations). Numerical results are reported in Supplementary Table 10.

To better understand the improvement of PolyFun + SuSiE over SuSiE, we examined the 121 loci where PolyFun + SuSiE identified a fine-mapped common SNP (PIP>0.95) but SuSiE did not (PIP<0.5 for all SNPs within 1Mb) (Figure 5 and Supplementary Table 16). In each case, functional annotations prioritized one SNP out of several candidates, greatly improving fine-mapping resolution. Examples included (a) height, for which rs288326, a non-synonymous SNP, had PIP=0.96 for PolyFun + SuSiE vs. PIP=0.35 for SuSiE (Figure 5a); (b) BMI, for which rs12330631, a conserved SNP, had PIP=0.96 for PolyFun + SuSiE vs. PIP=0.25 for SuSiE (Figure 5b); (c) red blood cell count, for which rs80207740, a promoter SNP, had PIP=0.97 for PolyFun + SuSiE vs. PIP=0.28 for SuSiE (Figure 5c); and (d) platelet count, for which rs2270894, a promoter SNP, had PIP=0.96 for PolyFun + SuSiE vs. PIP=0.19 for SuSiE (Figure 5d). Results for all 121 loci are reported in Supplementary Table 16.

**Figure 5:**
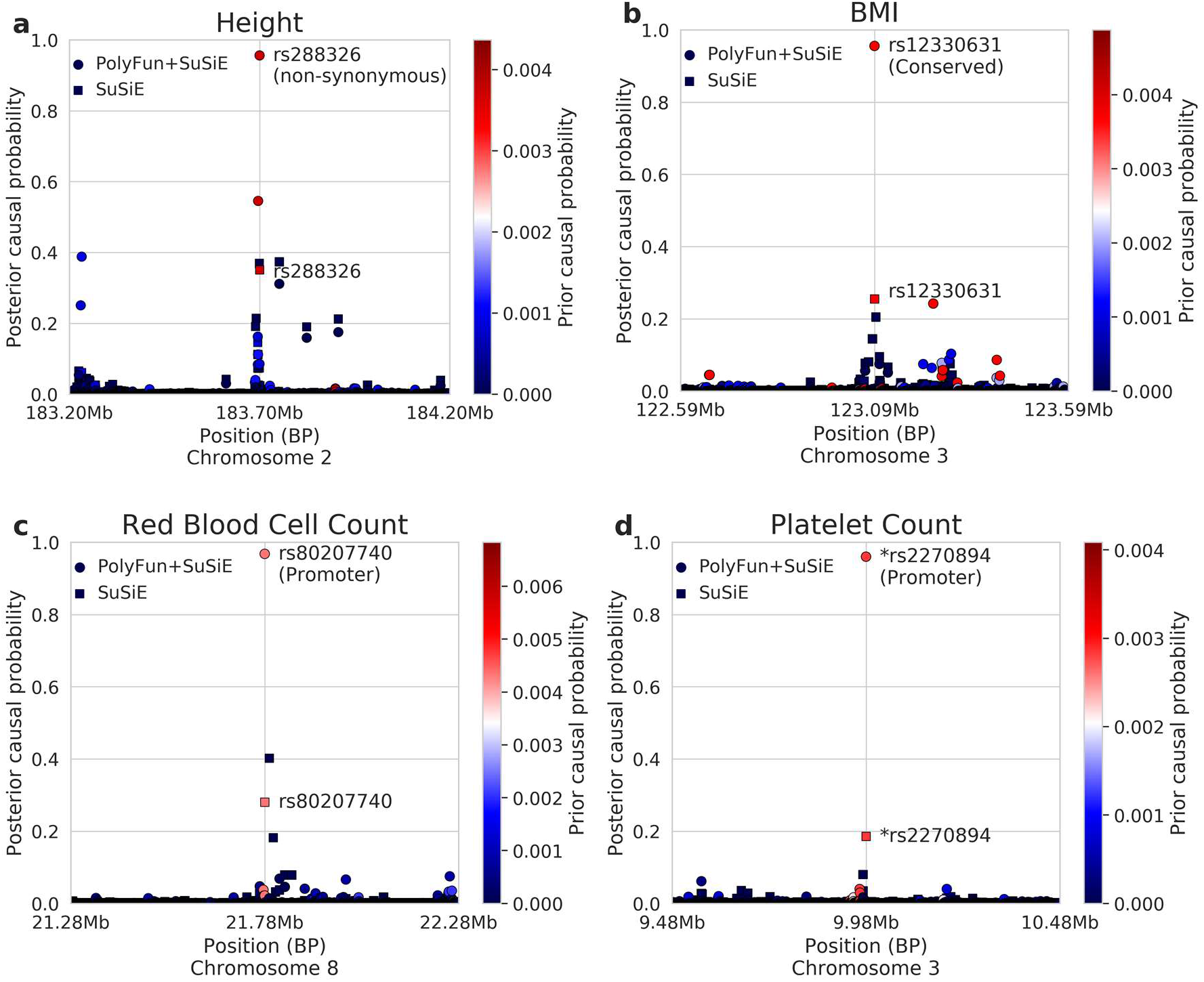
Examples of the advantages of functionally-informed fine-mapping for UK Biobank traits. We report four examples where PolyFun + SuSiE identified a fine-mapped common SNP (PIP>0.95) but SuSiE did not (PIP<0.5 for all SNPs within 1Mb). Circles denote PolyFun + SuSiE PIPs and squares denote SuSiE PIPs. SNPs are shaded according to their prior causal probabilities estimated by PolyFun. The top PolyFun + SuSiE SNP is labeled (next to its PolyFun + SuSiE PIP and its SuSiE PIP). The annotation of each top PolyFun + SuSiE SNP that is most enriched among SuSiE PIP>0.95 SNPs (Methods) is reported in parentheses below its label. Asterisks denote lead GWAS SNPs. Numerical results are reported in Supplementary Table 16.

We validated the motivation for performing functionally-informed fine-mapping by verifying that fine-mapped SNPs are enriched for functional annotations, as previously shown for autoimmune diseases^7,8,10^ and blood traits^12^. We used non-functionally-informed SuSiE in this analysis to avoid biasing the results. For each of 50 main binary annotations from the baseline-LF model^25^, for various PIP ranges, we computed the functional enrichment of fine-mapped common SNPs in the PIP range, defined as the proportion of common SNPs in the PIP range lying in the annotation divided by the proportion of genome-wide common SNPs lying in the annotation, and meta-analyzed the results across genetically uncorrelated traits (Methods, Figure 6, Supplementary Table 17). PIP>0.95 SNPs were strongly and significantly enriched for non-synonymous SNPs (51x enrichment, P=6.8×10^−185^) and SNPs in conserved regions (16x enrichment, P< 10^−300^), significantly enriched for SNPs in various regulatory annotations (e.g. promoter-ExAC and H3K4me3), and significantly depleted for SNPs in repressed regions, consistent with previous literature on functional enrichment of fine-mapped SNPs^7,8,10–12^ and disease heritability^17,25,26,45^. We observed qualitatively similar but weaker enrichments at lower PIP ranges (Figure 6, Supplementary Table 17).

**Figure 6:**
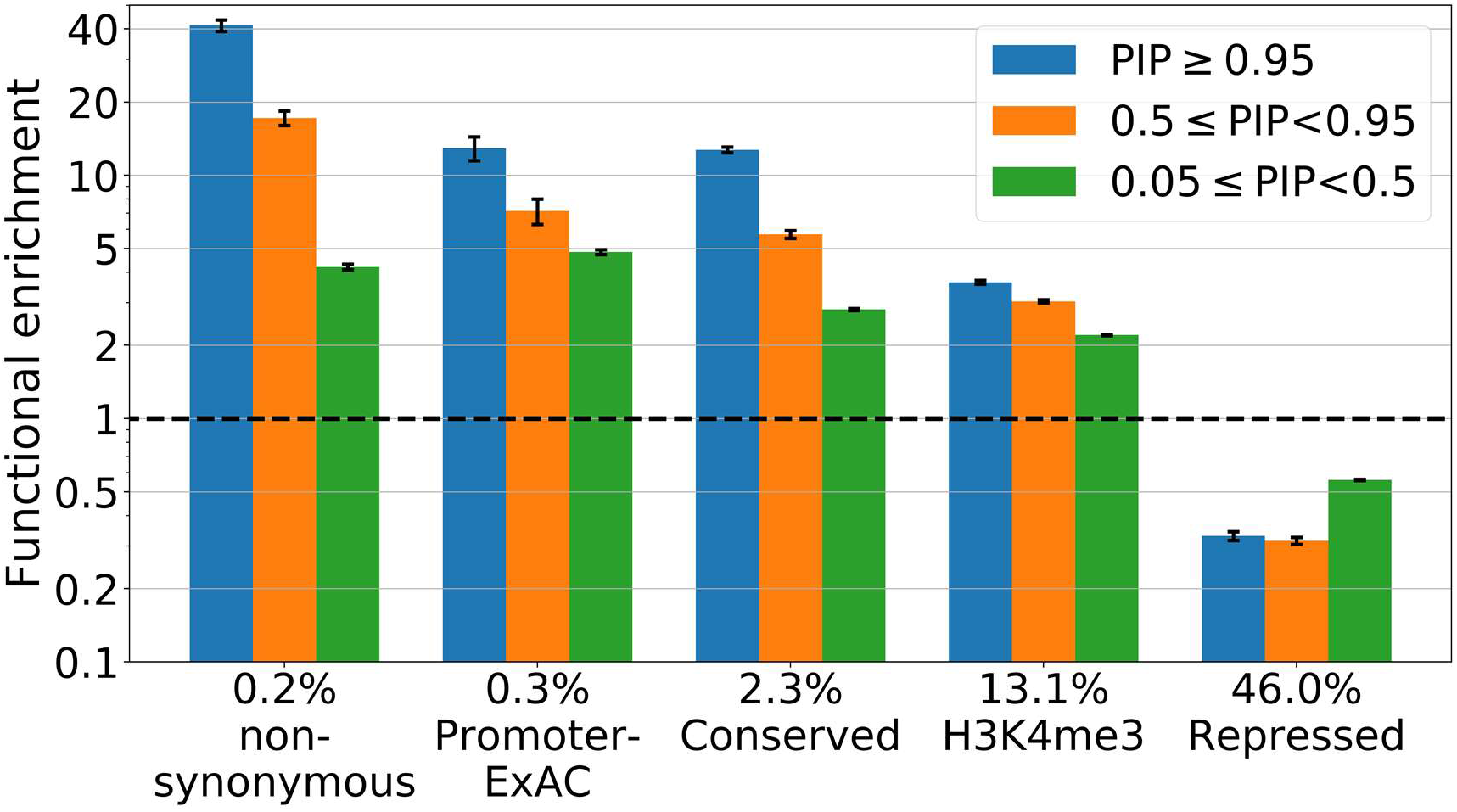
Functional enrichment of SuSiE fine-mapped common SNPs for UK Biobank traits. We report the functional enrichment of fine-mapped common SNPs (defined as the proportion of common SNPs in a PIP range lying in an annotation divided by the proportion of genome-wide common SNPs lying in the annotation) for 5 selected binary annotations, meta-analyzed across 14 genetically uncorrelated UK Biobank traits with ≥10 PIP>0.95 SNPs (log scale). The proportion of common SNPs lying in each binary annotation is reported above its name. The horizontal dashed line denotes no enrichment. Error bars denote standard errors. Numerical results for all 50 main binary annotations and all traits are reported in Supplementary Table 17.

We compared our fine-mapping results to two previous studies. First, we compared our results to ref. ^12^, which performed non-functionally informed fine-mapping (using a previous version of FINEMAP^23^) for the 9 blood cell traits using a subset of approximately 115K of the individuals included in our analyses. PolyFun + SuSiE identified 1,268 PIP>0.95 SNP-trait pairs (4.4× more than ref. ^12^; 289 PIP>0.95 SNP-trait pairs), 146 of which were shared (PIP>0.95) across the two studies, including all four SNPs that were functionally validated via luciferase reporter assays in ref. ^12^ (PIP>0.999 for all four SNPs; Methods, Supplementary Table 18-20). Sample size was the most important difference between the two studies, and incorporation of functional priors was also important, as we determined that (non-functionally informed) SuSiE identified only 984 PIP>0.95 SNP-trait pairs (3.4x more than ref. ^12^), 146 of which were shared (Supplementary Tables 18-19). Surprisingly, many differences remained even after we restricted SuSiE to the same subset of 115K individuals analyzed in ref. ^12^ (242 PIP>0.95 SNP-trait pairs, 130 of which were shared; Supplementary Tables 18-19), possibly due to differences in the underlying methods, data preprocessing, or reference panel used to impute genotypes. Second, we compared our results to ref. ^7^, which performed non-functionally-informed fine-mapping for 7 of our traits (bone mineral density, fasting glucose, HDL cholesterol, LDL cholesterol, platelet count, red blood cell count, triglycerides), using a non-functionally informed method (PICS) and independent smaller data sets. PolyFun + SuSiE identified 727 PIP>0.95 SNP-trait pairs (35x more than ref. ^7^; 21 PIP>0.95 SNP-trait pairs; Supplementary Tables 21-22). 12 SNP-trait pairs had PIP>0.95 in both studies, implying a replication rate (in independent data) of 57% (12/21) in our study. We caution that the fine-mapping power of PolyFun + SuSiE is likely much lower than 57% in practice, because the 21 SNPs fine-mapped in ref. ^7^ are likely to have larger effect sizes than most causal SNPs. We did not compare our results to refs. ^8,10,11^ because those studies analyzed disease traits, for which the number of cases in the UK Biobank is relatively low.

We performed 5 secondary analyses. First, we verified the robustness of our fine-mapping results by repeating the analysis using data from only the UK Biobank interim release (average *N*=107K) for the five traits with highest fine-mapping power (Methods): height, platelet count, bone mineral density, red blood cell count, and lymphocyte count. PolyFun + SuSiE identified >45% more PIP>0.95 SNPs vs. SuSiE overall, and >33% more PIP>0.95 SNPs vs. SuSiE among SNPs having PIP>0.95 in the full *N*=337K SuSiE results, with a similar rate of replication in the full *N*=337K SuSiE results (Supplementary Table 23). Second, in the *N*=337K analysis, we determined that the union of PolyFun + SuSiE 95% credible sets (defined as sets with probability >0.95 of including ≥1 causal SNP^22^; see above) was 23% smaller than the union of SuSiE 95% credible sets (median reduction) (Supplementary Table 24), consistent with simulations (Supplementary Table 4). Third, we searched for pairs of fine-mapped SNPs within 1Mb of each other where exactly one of the SNPs is coding, which can aid in linking regulatory variants to target genes^46–48^, and identified 490 such pairs (Supplementary Table 25). Fourth, we compared the magnitude of the posterior mean and posterior standard deviation of causal effect sizes, which can inform the applicability of fine-mapping results to polygenic risk scores. Posterior means were 3.6x smaller than posterior standard deviations (median ratio) for PIP>0.05 SNPs but 6.9x larger for PIP>0.95 SNPs, for which causal effect sizes are estimated with high accuracy (Supplementary Table 10). Fifth, we verified that the functional enrichments of fine-mapped SNPs remained qualitatively similar when using PolyFun + SuSiE instead of SuSiE (Supplementary Figure 1, Supplementary Table 26) and when defined using all MAF≥0.001 SNPs (Supplementary Figure 2, Supplementary Table 27) or only low-frequency and rare SNPs (0.05>MAF≥0.001) (Supplementary Figure 3, Supplementary Table 28).

In summary, we leveraged the improved power of PolyFun + SuSiE to robustly identify thousands of fine-mapped SNPs, providing a rich set of potential candidates for functional follow-up. Our results further indicate pervasive pleiotropy, with many SNPs fine-mapped for two or more genetically uncorrelated traits.

### Functionally-informed polygenic localization of 49 complex traits in the UK Biobank

PIP>0.95 SNPs tag a large proportion of the SNP-heritability 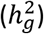 tagged by lead GWAS SNPs (median proportion=42%) but a small proportion of total genome-wide 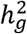 (median proportion=19%) (Figure 3b), implying that they causally explain a small proportion of 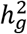. We thus propose *polygenic localization*, whose aim is to identify a minimal set of common SNPs causally explaining a specified proportion of common SNP heritability. A key difference between polygenic localization and previous studies of polygenicity^29,49–52^ is that polygenic localization aims to *identify* (not just characterize) such SNPs.

Given a ranking of SNPs by posterior per-SNP heritability (i.e., the posterior mean of their squared effect size; see Methods), we define *M*_50%_ as the size of the smallest set of top-ranked common SNPs causally explaining 50% of common SNP heritability (resp. *M*_*p*_ for proportion *p* of common SNP heritability). We estimate *M*_50%_ (resp. *M*_*p*_) by (1) partitioning SNPs into 50 ranked bins of similar posterior per-SNP heritability estimates from PolyFun + SuSiE and stratifying the lowest-heritability bin into 10 equally-sized MAF bins, yielding 59 bins; (2) running S-LDSC using a different set of samples to re-estimate the average per-SNP heritability in each bin; and (3) computing the number of top-ranked common SNPs (with respect to the original ranking) whose estimated per-SNP heritabilities (from step 2) sum up to 50% (resp. the proportion *p*) of the total estimated SNP-heritability. We refer to this method as PolyLoc. The analysis of new samples in step 2 of PolyLoc prevents winner’s curse; although PolyFun + SuSiE is robust to winner’s curse, PolyLoc would be susceptible to winner’s curse if it reused the data analyzed by PolyFun + SuSiE. We note that *M*_50%_ relies on an empirical ranking and is thus larger than the size of the smallest set of SNPs causally explaining 50% of common SNP heritability, denoted as 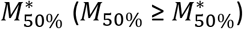. We further note that PolyLoc will yield robust estimates of *M*_50%_ if S-LDSC yields robust estimates of the SNP-heritability causally explained by each bin. Although S-LDSC has previously been shown to produce robust estimates^17,25–27^, we performed extensive simulations to confirm that PolyLoc produced robust estimates of *M*_50%_ (Methods, Supplementary Tables 32-33). Further details of PolyLoc are provided in the Methods section; we have released open source software implementing PolyLoc (see URLs).

We applied PolyLoc to the 49 complex traits from the UK Biobank (Supplementary Table 8). We ranked SNPs using *N*=337K unrelated British ancestry samples (steps 1-2) and re-estimated average per-SNP heritabilities in each of 59 SNP bins using S-LDSC applied to *N*=122K European-ancestry UK Biobank samples that were not included in the *N*=337K set (step 3). Estimates of *M*_50%_ ranged widely from 28 (hair color) to 3.4K (height) to 2 million (number of children; Figure 7, Supplementary Table 29). The median estimate of *M*_50%_ across 16 genetically uncorrelated traits was 8.9K; the median estimate of *M*_5%_ was 8; and the median estimate of *M*_95%_ was 4.4 million (of 7.0 million total common SNPs) (Supplementary Table 29). We verified that *M*_50%_ estimates were strongly correlated with estimates of the effective number of independently associated SNPs (*M_e_*; ref. ^29^), with log-scale *r*=0.92 (P=3.1×10^−6^) across 14 genetically uncorrelated traits with published *M_e_* estimates (Supplementary Table 30). Pigmentation traits were the least polygenic traits while number of children was the most polygenic trait, having *M*_50%_ 3.7x larger than the second most polygenic of the 16 independent traits (chronotype, having *M*_50%_=553K), consistent with ref. ^29^. As noted above, PolyLoc provides the SNP sets underlying its estimates, unlike ref. ^29^.

**Figure 7:**
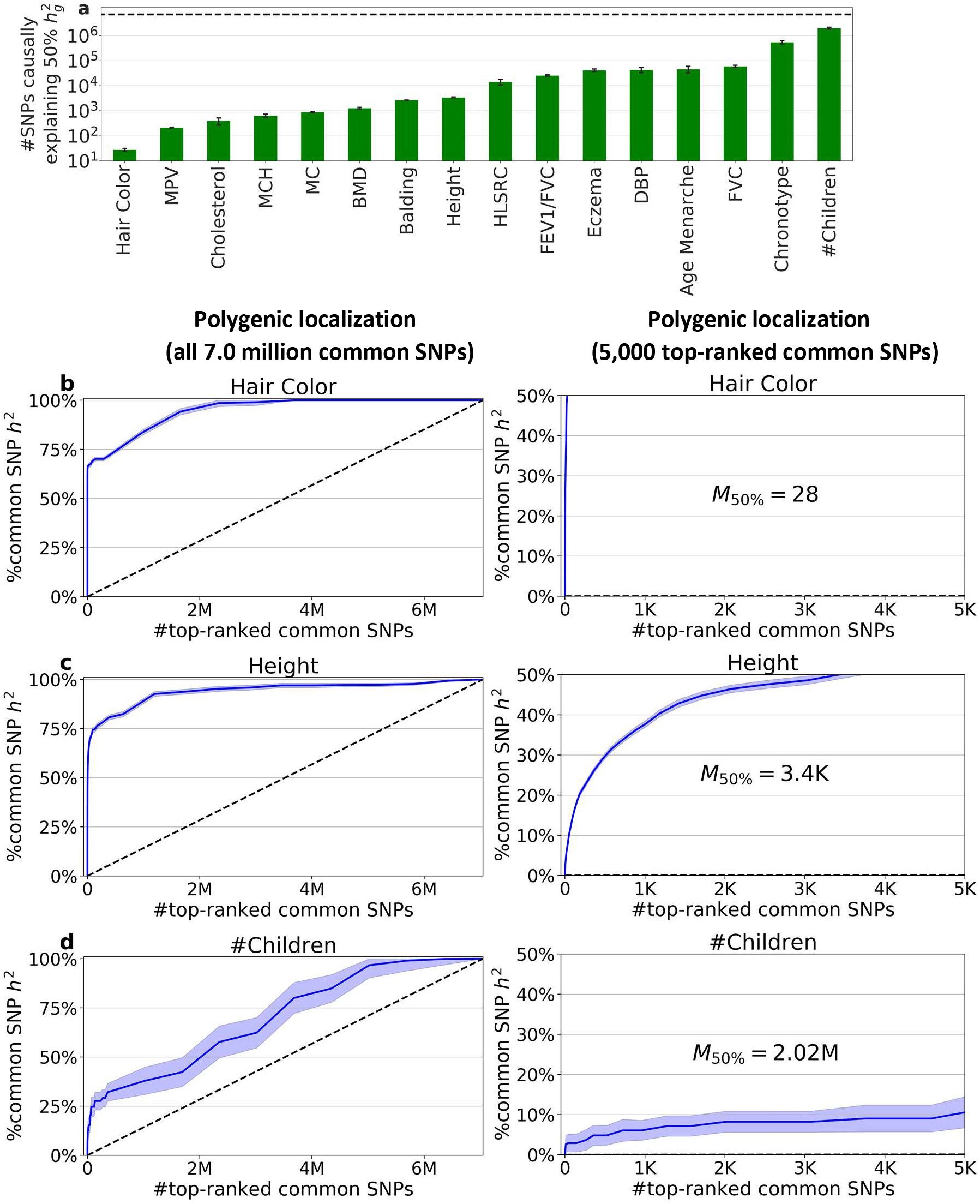
Polygenic localization results for UK Biobank traits. (**a**) *M*_50%_ estimates across 16 genetically uncorrelated traits. For each trait, we report the number of top-ranked common SNPs (using PolyFun + SuSiE posterior per-SNP heritability estimates for ranking) causally explaining 50% of common SNP heritability, and its standard error (log scale). The horizontal dashed line denotes the total number of common SNPs in the analysis (7.0 million). (**b-d**) The proportion of common SNP heritability of (b) hair color, (c) height, and (d) number of children explained by different numbers of top-ranked SNPs, for all 7.0 million common SNPs (left) and the 5,000 top-ranked common (right). Purple shading denotes standard errors. Dashed black lines denote a null model with a constant per-SNP heritability. We also report the number of top-ranked SNPs causally explaining 50% of common SNP heritability, denoted *M*_50%_. Discontinuities in the slope indicate transitions between SNP bins. Numerical results for all 49 UK Biobank traits are reported in Supplementary Table 29.

We performed 6 secondary analyses. First, we used posterior per-SNP heritability estimates from SuSiE (instead of PolyFun + SuSiE) to partition SNPs into heritability bins, obtaining *M*_50%_ estimates that were 1.4x larger (median ratio; Supplementary Table 31). This result is consistent with our results showing that PolyFun + SuSiE identifies more PIP>0.95, PIP>0.5 and PIP>0.05 SNPs than SuSiE (Supplementary Table 9), and further illustrates the improved fine-mapping resolution of PolyFun + SuSiE vs. SuSiE. Second, we applied PolyLoc to prior estimates of per-SNP heritability computed by S-LDSC based only on functional annotations (including MAF bins; Methods) and obtained *M*_50%_ estimates that were overwhelmingly larger, emphasizing the value of posterior estimates (Supplementary Table 31). Third, we modified PolyLoc by applying step (3) of PolyLoc to the same *N*=337K samples analyzed by PolyFun + SuSiE. We obtained *M*_50%_ estimates that were drastically different, a consequence of winner’s curse (Supplementary Table 31). Fourth, we verified that PolyLoc results were not sensitive to the number of heritability bins (Supplementary Table 31). Fifth, we determined that polygenic localization estimates were slightly larger when not stratifying the lowest-heritability bin into 10 MAF bins (Supplementary Table 31). Sixth, We determined that *M*_50%_ estimates using all MAF≥0.001 SNPs were 1.4x larger (vs. 2.8x larger SNP set) compared to analyses of common SNPs, confirming that rare and low-frequency SNPs are depleted for high-heritability SNPs^26,49,53^ (Supplementary Table 31).

Our results demonstrate that half of the common SNP heritability of complex traits is causally explained by typically thousands of SNPs (median *M*_50%_=8.9K), and the remaining heritability is spread across an extremely large number of extremely weak-effect SNPs (median *M*_95%_=4.4 million), consistent with extremely polygenic but heavy-tailed trait architectures^1,29,49,50,54–58^.

## Discussion

We have introduced PolyFun, a framework that improves fine-mapping by prioritizing variants that are a-priori more likely to be causal based on their functional annotations. Across 49 UK Biobank traits, PolyFun + SuSiE confidently fine-mapped 3,025 SNP-trait pairs (PIP >0.95), a 32% increase over non-functionally informed SuSiE. The fine-mapped SNPs span 20% of GWAS loci but explain 42% of lead GWAS SNP-heritability, as fine-mapping power is higher at loci with larger causal effects. 223 of the fine-mapped SNPs were fine-mapped for multiple genetically uncorrelated traits, indicating pervasive pleiotropy. PolyFun improves our ability to scale the identification of causal variants underlying association signals, a primary challenge in genetics research^2^. We further leveraged the results of PolyFun to perform polygenic localization by constructing minimal SNP sets causally explaining a given proportion of common SNP heritability, demonstrating that 50% of common SNP heritability can be explained by sets ranging in size from 28 (hair color) to 3,400 (height) to 2 million (number of children). We have publicly released the PIPs and the prior and posterior means and variances of effect sizes for all SNPs and traits analyzed (see URLs).

We recommend applying PolyFun using in-sample LD from the GWAS target sample, assuming 10 causal SNPs per locus; we have facilitated this option for UK Biobank researchers by publicly releasing summary LD information for British-ancestry UK Biobank samples as part of this study (see URLs). As a second-best option we recommend applying PolyFun using LD-reference panel from the same population as the target sample that spans at least 10% of the target sample size while assuming 10 causal SNPs per locus. However, we caution that even subtle population differences may lead to false positive results. In the absence of a reference panel from the exact same population as the target sample that spans >10% of the target sample size, we recommend applying PolyFun without using an LD reference panel by restricting PolyFun to assume a single causal SNP per locus.

PolyFun allows for fine-mapping all genome-wide loci, instead of just genome-wide significant loci. For each locus analyzed, PolyFun fine-maps all signals in the locus jointly to maximize power (instead of partitioning the locus into independent signals^59^ and fine-mapping those signals separately). Specifically, previous studies have shown that fine-mapping methods that partition a locus into independent signals do not make full use of available information in the case of multiple causal variants in partial LD^5,30^. Researchers wishing to use PolyFun for a partitioned analysis may still do so by first partitioning a locus into multiple signals using a separate tool (e.g. GCTA-COJO^59^) and then applying PolyFun to each signal separately, restricting PolyFun to assume a single causal SNP per signal.

Our results provide several opportunities for future work. First, the fine-mapped SNPs that we have identified can be prioritized for functional follow-up. Second, fine-mapping results (posterior mean effect sizes) can be used to compute polygenic risk scores^60^. This may be especially useful for cross-population prediction, which is sensitive to misspecification of causal SNPs due to LD differences between populations^61,62^. Third, the proximal pairs of coding and non-coding fine-mapped SNPs that we identified (Supplementary Table 25) may aid efforts to link SNPs to genes^46–48^. Fourth, SNPs that were fine-mapped for multiple genetically uncorrelated traits may shed light on shared biological pathways^63^. Fifth, sets of SNPs causally explaining 50% of common SNP heritability can potentially be used for gene and pathway enrichment analysis^64,65^. Finally, PolyFun can incorporate additional functional annotations at negligible additional computational cost, motivating further efforts to identify conditionally informative annotations.

Our work has several limitations. First, our PIP≥0.95 FDR estimates for PolyFun and for other methods are conservative. We have demonstrated that an alternative data-driven estimator is anti-conservative and hence not recommended (Methods), demonstrating the challenges of exact calibration in fine-mapping. We are not currently aware of any way to produce an FDR estimate for imputed data that is not conservative or anti-conservative (in particular, we note that the recently proposed method KnockOffZoom^66^ provably controls FDR in the case of direct genotype data, but cannot provably control FDR in the case of imputed data; it may be possible to combine PolyFun and KnockOffZoom (PolyFun + KnockOffZoom) by applying KnockOffZoom to priors from PolyFun (analogous to PolyFun + SuSiE), a direction for future work). Second, subtle population stratification may lead to spurious fine-mapping results, arising from spurious association results^67^. However, our fine-mapped SNPs are concentrated in associated loci with larger estimated effects, which are relatively less likely to be spurious. Third, we did not compare PolyFun to the functionally informed fine-mapping method applied in ref. ^10^ (an extension of fGWAS^21^), which was not made publicly available. However, that method can only incorporate a limited number of functional annotations (e.g. <15 in ref. ^10^) and uses stepwise conditional fine-mapping, which has been shown to be susceptible to spurious findings^30^. Fourth, PolyFun + SuSiE assumes 10 causal SNPs per locus (Table 1). This choice has been shown to have minimal impact on SuSiE results^22^, but it may still be advantageous to assess the number of causal SNPs per locus in a data-driven manner. Fifth, PolyFun (and SuSiE) were designed for quantitative phenotypes but can also be applied to binary phenotypes; alternative methods designed for binary phenotypes may further increase power. Sixth, we restricted our analyses to MAF>0.001 SNPs, because SNPs with lower MAFs are often not well-imputed^37^. Future studies with whole genome sequencing data could potentially fine-map MAF≤0.001 SNPs, but the performance of PolyFun + SuSiE on sequencing data has not been investigated. Seventh, we performed fine-mapping using 3Mb windows, but in rare cases causal SNPs might be in LD with associated SNPs that are outside these windows. Eighth, application of PolyLoc requires analyzing samples distinct from the samples analyzed by PolyFun to avoid winner’s curse. Researchers with access to individual-level genetic data can partition the samples as we have done (we recommend using approximately 75% of the data for fine-mapping and 25% for polygenic localization). A potential alternative is to apply methods to alleviate bias due to winner’s curse^68,69^. We emphasize that not correcting for winner’s curse can lead to extremely biased estimates (Supplementary Table 31). Ninth, PolyLoc provides an upper bound on the proportion of SNPs causally explaining a given proportion of SNP-heritability and is thus conservative. Finally, multi-ethnic fine-mapping^70^ and incorporation of tissue-specific functional annotations^9,13,15,17^ may further increase fine-mapping power. Incorporating these into our fine-mapping framework is an avenue for future work.

## Methods

### PolyFun fine-mapping method

PolyFun performs functionally-informed fine-mapping by first estimating prior causal probabilities for all SNPs and then applying fine-mapping methods such as SuSiE^22^ or FINEMAP^23,24^ with these prior causal probabilities. Here we describe estimation of the prior causal probabilities.

We model standardized phenotypes *y* using the linear model *y* = ∑_*i*_ *x*_*i*_*β*_*i*_ + *ϵ*, where *x*_*i*_ denotes standardized SNP genotypes, *β*_*i*_ denotes effect size, and *ϵ* is a residual term. We use a point-normal model for *β*_*i*_:

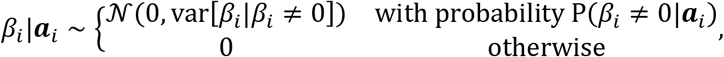

where ***α***_*i*_ are the functional annotations of SNP *i*, *P*(*β*_*i*_ ≠ 0|***α***_*i*_) is its prior causal probability, and var[*β*_*i*_|*β*_*i*_ ≠ 0] is its causal variance, which we assume is independent of ***α***_*i*_. This assumption is motivated by our recent work showing that functional enrichment is primarily due to differences in polygenicity rather than differences in effect-size magnitude, which is constrained by negative selection^29^.

The key quantity that PolyFun uses to estimate prior causal probabilities is the per-SNP heritability of SNP *i*, var[*β*_*i*_|***α***_*i*_] (we refer to this quantity as per-SNP heritability because the total SNP-heritability var[∑_*i*_ *x*_*i*_*β*_*i*_ |***α***] is equal to ∑_*i*_ var[*β*_*i*_|***α***_*i*_], assuming that causal SNP effects have zero mean and are uncorrelated with other SNP effects and with other SNPs conditional on ***α***). PolyFun relates the prior causal probability P(*β*_*i*_ ≠ 0|***α***_*i*_) to the per-SNP heritability var[*β*_*i*_|***α***_*i*_] via the law of total variance:

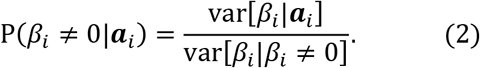

Equation 1 in the main text follows since P(*β*_*i*_ ≠ 0|***α***_*i*_) is proportional to var[*β*_*i*_|***α***_*i*_] with the proportionality factor 1/var[*β*_*i*_|*β*_*i*_ ≠ 0].

To derive Equation 2 we define the causality indicator 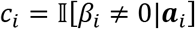 and apply the law of total variance to var[*β*_*i*_|***α***_*i*_]:

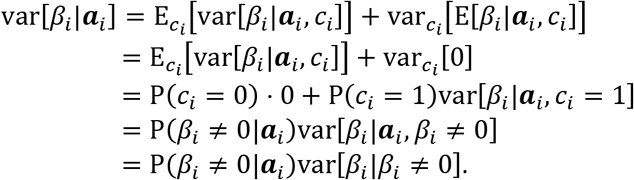

The last equality holds because we assume that the causal effect size variance is independent of functional annotations, as explained above.

PolyFun avoids directly estimating the proportionality factor 1/var[*β*_*i*_|*β*_*i*_ ≠ 0] by constraining the prior causal probabilities P(*β*_*i*_ ≠ 0|***α***_*i*_) in each tested locus to sum to 1.0. This constraints implies that each locus is a-priori expected to harbor one causal SNPs, consistent with previous fine-mapping methods^5,6,23^ (we note that this constraint is ignored by PolyFun + SuSiE, because it is invariant to scaling of prior causal probabilities). Hence, the main challenge is estimating the per-SNP heritabilities var[*β*_*i*_|***α***_*i*_].

To estimate var[*β*_*i*_|***α***_***i***_], PolyFun incorporates a regularized extension of S-LDSC with the baseline-LF model^17,25–27^, which we extend to a new version 2.2.UKB (Supplementary Table 1, URLs, see below). S-LDSC uses the linear model 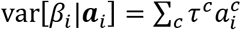 and jointly estimates all *τ*^*c*^ parameters by minimizing the term 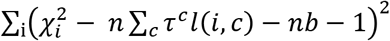, where *c* are functional annotations, *τ*^*c*^ is the coefficient of annotation *c*, 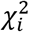 is the *χ*^2^ statistic of SNP *i*, *n* is the sample size, *b* measures the contribution of confounding biases, and 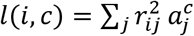.

While S-LDSC produces robust estimates of functional enrichment, it has two limitations in estimating var[*β*_*i*_|***α***_*i*_]: (i) these estimates can have large standard errors in the presence of many annotations, and (ii) the model may not be robust to model misspecification. To address the first limitation, PolyFun incorporates an L2-regularized extension of S-LDSC. To address the second limitation, PolyFun employs special procedures to ensure robustness to model misspecification. The key idea is to approximate arbitrary complex functional forms of var[*β*_*i*_|***α***_*i*_] via a piecewise-constant function. To do this, PolyFun partitions SNPs with similar estimated values of var[*β*_*i*_|***α***_*i*_] (estimated via a possibly misspecified model) into non-overlapping bins; estimates the SNP-heritability causally explained by each bin *b*; and specifies var[*β*_*i*_|***α***_*i*_] for SNPs in bin *b* as the SNP-heritability causally explained by bin *b* divided by the number of SNPs in bin *b*. PolyFun avoids winner’s curse by using different data for partitioning SNPs and for per-bin heritability estimation.

In detail, PolyFun robustly specifies prior causal probabilities for all SNPs on a target locus on a corresponding odd (resp. even) target chromosome via the following procedure:

1. Estimate annotation coefficients 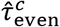 and intercepts 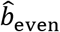 using only SNPs in even chromosomes via an L2-regularied extension of S-LDSC that minimizes 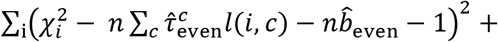 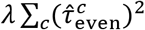 (resp. using 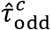 and 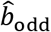). We select the regularization strength *λ* from a geometrically-spaced grid of 100 values ranging from 10^−8^ to 100, selecting the one that minimizes the average out-of-chromosome error 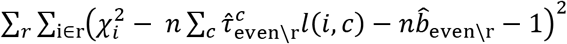, where *r* iterates over even (resp. even) chromosomes, and 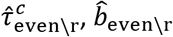 are the S-LDSC *τ* and *b* estimates, respectively, when applied to all SNPs on even chromosomes except for chromosome *r* (resp. for odd chromosomes).
2. Compute per-SNP heritabilities 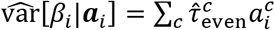 for each SNP *i* in an odd chromosome (resp. 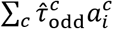).
3. Partition all SNPs into 20 bins with similar values of 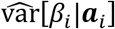 using the Ckmedian.1d.dp method^71^. This method partitions SNPs into 20 maximally homogenous bins such that the average distance of 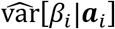 to the median 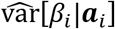 of the bin of SNP *i* is minimized. We emphasize that even though this step uses functional annotations data of the target chromosome it does not use the summary statistics of SNPs in the target chromosome, which ensures robustness to winner’s curse.
4. Apply S-LDSC with non-negativity constraints to estimate per-SNP heritabilities in each of the 20 bins of all SNPs in odd (resp. even) chromosomes except for the target chromosome *r* (to avoid using the same data that will be used in fine-mapping), denoted 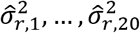. Afterwards, regularize the estimates by setting all values smaller than 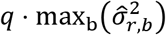 to 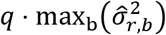, using *q* = 1/100 by default, and rescaling the 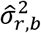 estimates to have the same sum (over all genome-wide SNPs) as before. The regularization prevents SNPs from a having a zero per-SNP probability, which would exclude them from fine-mapping. We did not apply L2-regularization in this step because we require approximately unbiased estimates, and because standard errors are relatively small in the presence of a small number of non-overlapping annotations.
5. Specify a prior causal probability proportional to 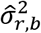 to each SNP that is in bin *b* and that resides in a target locus in chromosome *r*, such that the prior causal probabilities in the target locus sum to one.

PolyFun uses version 2.2.UKB of the baseline-LF model, which differs from the original baseline-LF model^26^ by including MAF≥0.001 SNPs and several new annotations, and omitting annotations that could not be easily extended to account for MAF<0.005 SNPs (Supplementary Table 1). Briefly, we use 187 overlapping functional annotations, including 10 common MAF bins (MAF≥0.05); 10 low-frequency MAF bins (0.05>MAF≥0.001); 6 LD-related annotations for common SNPs (levels of LD, predicted allele age, recombination rate, nucleotide diversity, background selection statistic, CpG content); 5 LD-related annotations for low-frequency SNPs; 40 binary functional annotations for common SNPs; 7 continuous functional annotations for common SNPs; 40 binary functional annotations for low-frequency SNPs; 3 continuous functional annotations for low-frequency SNPs; and 66 annotations constructed via windows around other annotations^17^. We did not include a base annotation that includes all SNPs, because such an annotation is linearly dependent on all the MAF bins when S-LDSC uses the same set of SNPs to compute LD-scores and to estimate annotation coefficients.

### False discovery rate analysis

Our simulation studies indicate that the PIPs reported by PolyFun+SuSiE and PolyFun+FINEMAP are anti-conservative with respect to the expected FDR at PIP=0.95, defined as one minus the average PIP among PIP≥0.95 SNPs (lower dotted horizontal lines in Figures 1a-b). (For example, if all SNPs with PIP>0.95 have PIP=1.0 then the expected FDR is 0% rather than 5%, which would imply that none of the PIP>0.95 SNPs are false-positives.) To circumvent this problem, we propose a heuristic FDR given by one minus the PIP threshold (upper dashed horizontal lines in Figures 1a-b). This definition can be derived by upper bounding all PIPs to the PIP threshold and then computing the expected FDR. For example, for a PIP threshold of 0.95, we propose to treat all SNPs with PIP≥0.95 as if they had PIP=0.95. Our simulation studies indicate that this heuristic definition of FDR typically yields conservative FDR estimates.

We now explain the rationale behind our heuristic definition of FDR, assuming without loss of generality a PIP threshold of 0.95. We begin by writing down the definition of expected FDR under a Bayesian framework, given by 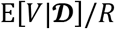, where *V* is the number of non-causal PIP>0.95 SNPs, *R* is the number of PIP>0.95 SNPs, and 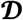 is the GWAS and LD data used for fine-mapping. Let us define *V* = ∑_*i*_ *I*_*i*_, where *i* iterates over PIP>0.95 SNPs, and *I*_*i*_ is an indicator that is equal to 0 if SNP *i* is causal and to 1 otherwise. Then:

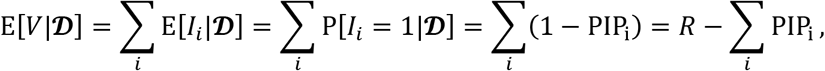

where PIP_*i*_ is the PIP of SNP *i*. We conclude that 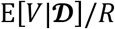 is given by one minus the average PIP among PIP>0.95 SNPs.

As mentioned above, estimates of expected FDR under this definition are anti-conservative. Although it is difficult to identify the reasons for this anti-conservativeness, possible reasons are (1) PolyFun + SuSiE and PolyFun + FINEAP compute approximate rather than exact PIPs; and (2) the prior causal probabilities we provided to these methods are imperfectly estimated due to having a finite sample size. To circumvent this problem we use the upper-bound 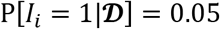. This modification is equivalent to treating all PIPs above 0.95 as if they were equal to 0.95. This heuristic definition of FDR typically yields conservative FDR estimates.

To gain a better understanding of these FDR estimates, we consider a hypothetical fine-mapping study with exactly 200 PIP≥0.95 SNPs, consisting of 40 PIP=0.96 SNPs, 40 PIP=0.97 SNPs, 40 PIP=0.98 SNPs, 40 PIP=0.99 SNPs, and 40 PIP=1.0 SNPs. Let us assume that 5 out of the 200 PIP≥0.95 SNPs are non-causal. The average PIP among the PIP≥0.95 SNPs is 0.98, yielding an expected FDR of 2%, but the true FDR is 2.5% (5/200 SNPs). Hence, the expected FDR is slightly anti-conservative (2.5% > 2%). To circumvent this problem, we transform the PIPs of all 200 PIP≥0.95 to 0.95. The average PIP among PIP≥0.95 SNPs is now 0.95, yielding a heuristic FDR of 5%. The true FDR remains 2.5% as before (5/200 SNPs). Hence the heuristic FDR is conservative (2.5% < 5%).

### Main fine-mapping simulations

We simulated summary statistics for 18,212,157 genotyped and imputed MAF≥0.001 autosomal SNPs with INFO score≥0.6 (including short indels, excluding three long-range LD regions; see below), using *N*=337,491 unrelated British-ancestry individuals from UK Biobank release 3. In most simulations we computed an effect variance *β*_*i*_ for every SNP *i* with annotations ***α***_*i*_ using the baseline-LF (version 2.2.UKB) model, 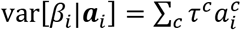, where *c* are annotations and *τ*^*c*^ estimates are taken from a fixed-effects meta-analysis of 16 well-powered genetically uncorrelated (|*r_g_*|<0.2) UK Biobank traits (age of menarche, BMI, balding, bone mineral density, eosinophil count, FEV1/FVC ratio, forced vital capacity, hair color, height, platelet count, red blood cell distribution width, red blood cell count, systolic blood pressure, tanning, waist-hip ratio adjusted for BMI, white blood count), scaled such that ∑_*i*_ var[*β*_*i*_|***α***_*i*_] is the same across all traits (Supplementary Table 3). In some simulations we generated values of var[*β*_*i*_|***α***_*i*_] under alternative functional architectures to evaluate the robustness of PolyFun to modeling misspecification (see below). Each SNP was set to be causal with probability proportional to var[*β*_*i*_|***α***_*i*_], such that the average causal probability was equal to the desired proportion of causal SNPs.

To facilitate the simulations we generated summary statistics directly, without first generating phenotypic values. This is mathematically equivalent to direct simulations but is substantially faster and allows modifying the residual variance for any desired sample size *n*. Specifically, we sampled a vector of marginal effect sizes ***α*** in each locus from 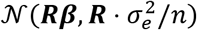 where ***R*** is a matrix of summary LD information (computed via LDstore^30^), ***β*** are effect sizes sampled iid from 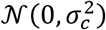 for causal SNPs (and set to 0 for non-causal SNPs), and 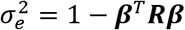 is the phenotypic variance not causally explained by SNPs in the locus^23,72^ (see below). The causal variance 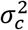 was set to 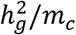 where 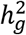 is the desired SNP heritability and *m*_*c*_ is the expected number of causal SNPs. We note that modifying *n* implies that ***R*** represents summary LD information from the population rather than from the sample.

We used the above procedure in two ways. First, we generated 100 sets of summary statistics in each of 10 3Mb loci in chromosome 1 for the fine-mapping experiments (selected by sorting all loci according to number of SNPs and selecting a uniformly-spaced set of loci spanning the two loci with the smallest and largest number of SNPs; Supplementary Table 2). We set 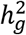 to the desired locus heritability and *m*_*c*_ to the desired number of causal SNPs in the locus, for a total of 1000 simulations (for simulations based on SuSiE and FINEMAP with default parameters) or 100 simulations (for all other simulations, due to computational cost) per unique combination of settings. Second, we generated 5 sets of genome-wide summary statistics for functional enrichment estimation, setting 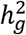 to the desired genome-wide SNP-heritability and *m*_*c*_ to the expected genome-wide number of causal SNPs. In the genome-wide simulations we generated a summary statistic *α*_*i*_ for each SNP based on the locus whose center was closest to the SNP among 2,763 overlapping 3Mb loci spanning all autosomal chromosomes (with a 1Mb spacing between the start points of consecutive loci and excluding three long-range LD regions: chr6 25.5M-33.5M, chr8 8M-12M, chr11 46M-57M; see below). In each experiment we randomly selected one of the 5 genome-wide sets of summary statistics and used it to estimate functional enrichment.

The default parameters for the locus-specific simulations were *m*_*c*_=10 and 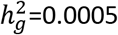 (broadly consistent with empirical data; average 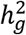 of a 3Mb locus across 16 genetically uncorrelated traits=0.0003). The default parameters for the genome-wide simulations were *m*_*c*_=91,700 (implying a proportion 0.5% of causal SNPs) and 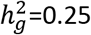 (consistent with real data results; Supplementary Table 14).

The most challenging aspect of the simulations is sampling vectors from 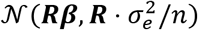, as it is both computationally intensive and technically complex due to singularity of ***R***. To circumvent these challenges we sampled 100 ***α***_*e*_ vector from *N*(**0**, ***R***) for each locus and stored them for future use. Afterwards, we generated summary statistics in each simulation via 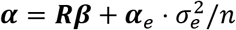, randomly choosing one of the 100 ***α***_*e*_ vectors. Because ***R*** was typically singular, we sampled ***α***_*e*_ approximately by (1) taking a random maximal subset of SNPs such that no pair of SNPs has |*r*|>0.99, using the maximal_independent_set procedure in the NetworkX package^73^, and constructing the corresponding submatrix ***R***_*s*_; (2) Finding the minimum value of *γ* such that (1 − *γ*)***R***_*s*_ + *γI* has a minimal eigenvalue >0.0001; (3) sampling ***α***_*e,s*_ from *N*(**0**, (1 − *γ*)***R***_*s*_ + *γI*); (4) sampling a value 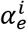 for each SNP *i* that was omitted in step 1 from a normal distribution conditional on 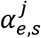 of the non-omitted SNP *j* having the strongest LD with SNP *i*; and (5) constructing ***α***_*e*_ by combining ***α***_*e,s*_ and all values of 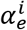.

After generating summary statistics, we first estimated prior causal probabilities for all SNPs as described in the PolyFun fine-mapping method subsection. We ran S-LDSC using LD scores computed from the summary LD information used to generate summary statistics (based on imputed SNP dosages rather than sequenced genotypes as in previous publications that used S-LDSC^25–27^, assigning to each SNP the LD score computed in the locus in which it was most central) to obtain sufficient coverage of low-frequency SNPs, which are underrepresented in small external reference panels.

We performed fine-mapping in each of the 10 selected 3Mb loci on chromosome 1 using methods based on SuSiE^22^, FINEMAP^23,24^, CAVIARBF^20^ and fastPAINTOR^19^. Following previous literature^12,30^ all methods used summary LD information computed via LDstore^30^ from the genotypes of the same 337,491 individuals used to generate summary statistics.

For fastPAINTOR-, fastPAINTOR, SuSiE, and PolyFun + SuSiE, we specified a causal effect size variance using an estimator that we developed based on a modified version of HESS^74^ rather than using the estimator implemented in these methods, because it improved false discovery rate and power in most simulation settings (Supplementary Table 4). Briefly, HESS estimates regional SNP-heritability via ***αR***^−1^***α*** − *m*/*n*, where ***α*** is a vector of marginal effect size estimates for *m* standardized SNPs, ***R*** is a matrix of summary LD information, and *n* is the sample size. We regularized this estimator (using summary LD information computed directly from the genotypes of the individuals used to generate summary statistics) by (1) excluding SNPs having 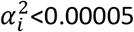 (i.e., having a negligible contribution to the estimator); and (2) selecting a random maximal subset of the remaining SNPs such that no pair of SNPs has |*r*|>0.99. We averaged this estimate across 100 different estimates per locus, each time selecting a different random subset via the maximal_independent_set procedure in the NetworkX package^73^. We then estimated the causal effect size variance as the HESS estimator divided by the assumed number of causal SNPs (using a value of 10 by default in this work). The division assumes that the correlation between causal SNPs is zero in expectation.

We now describe the parameters provided to each fine-mapping method.

We ran SuSiE 0.7.1.0487 with default values for all parameters except the following: (1) We used 10 causal SNPs per locus; and (2) we estimated a per-locus causal effect size variance (the scaled_prior_variance parameter) via our modified HESS approach. We specified prior causal probabilities via the prior_weights parameter. We modified the SuSiE source code to avoid performing the LD matrix diagnostics (positive-definiteness and symmetry) because they greatly increased memory consumption.

We ran FINEMAP 1.3.1.b (a new version of FINEMAP that we introduce here that incorporates prior causal probabilities) with a maximum of 10 causal SNPs per locus and with default settings for all other parameters. We specified prior causal probabilities via the –prior-snps argument.

We ran CAVIARBF 0.2.1 with an AIC-based parameter selection, using ridge regression with regularization parameter *λ* selected from {2^−10^, 2^−5^, 2^−2.5^, 2^0^, 2^2.5^, 2^5^, 100, 1000, 10000, 100000}, with a single locus and with up to either 1 or 2 causal SNPs per locus, owing to computational limitations.

We ran fastPAINTOR 3.1 in MCMC mode. We specified a per-locus causal effect size variance (specified via the -variance argument) using our modified HESS approach (as in PolyFun + SuSiE). We avoided truncation of the LD matrix (using prop_ld_eigenvalues=1.0) because we used in-sample summary LD information. As fastPAINTOR is generally not designed to work with >10 annotations^18,19^ (and was too slow in our simulations to estimate the significance of each annotation and include only conditionally significant annotations as done in ref. ^18^), we selected a subset of 10 highly informative annotations by (1) scoring each annotation based on its average contribution to effect variance 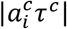 across all SNPs, using the true *τ*^*c*^ of the generative model; (2) iteratively selecting top-ranked annotations such that no annotation has correlation >0.3 (in absolute value) with a previously selected annotation, until selecting 10 annotations. We determined that 10 annotations yielded approximately optimal power while maintaining correct calibration (Supplementary Table 4).

For each PIP threshold, we conservatively estimated false discovery rates by setting all PIPs greater than the threshold to the threshold, yielding a uniform false-discovery threshold. This differs from exact false-discovery thresholds, defined as one minus the average PIP across all SNPs with PIP greater than the PIP threshold (e.g., if all SNPs with PIP>0.95 have PIP=1 then the expected false-discovery rate is 0% rather than 5%). We evaluated the results with respect to exact false-discovery thresholds in secondary analyses.

In a subset of the simulations we evaluated two alternative functional architectures: (1) A multiplicative functional architecture defined by 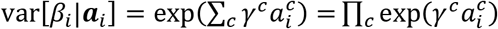, where *γ*^*c*^ is the coefficient of annotation *c*; and (2) a sub-additive functional architecture defined by 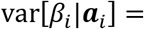 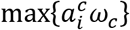, where *ω*_*c*_ is the coefficient of annotation *c*. To obtain realistic functional architectures, we fitted the coefficients of these two models to obtain a distribution of var[*β*_*i*_|***α***_*i*_] that is roughly similar to the distribution obtained under the standard S-LDSC with the baseline-LF model (with meta-analyzed linear annotation coefficients *τ*^*c*^ ; see above). In the multiplicative functional architecture, we fitted *γ*^*c*^ by approximately minimizing the mean squared distance between 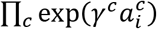 and 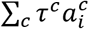 (via a linear regression with 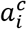 as explanatory variables and 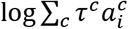 as the outcome, setting all estimates smaller than the median of the non-negative values of 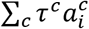 to the median to prevent the regression from being dominated by SNPs with low values of 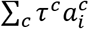). In the sub-additive functional architecture, we partitioned annotations into six groups (non-synonymous, coding, conserved, promoter or enhancer, histone marks, repressed, others) and associated all annotations in each group with the same coefficient *ω*_*c*_, such that the mean squared distance between 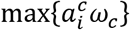 and 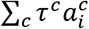 is minimized.

### Simulations with mismatched reference LD

We performed extensive simulations to assess how fine-mapping performance is impacted by LD mismatch between the target sample and the LD reference panel, focusing on (1) non-overlapping target and reference samples; (2) sample sizes of the target sample and reference panel; (3) differences in ancestry; (4) presence of related individuals in the target sample; and (5) SNPs available for analysis in the target sample and reference panel (Table 3).

Our mismatched reference LD simulations differed from our main simulations in several ways: (i) we generated summary statistics using up to *N*=44K unrelated (or related) European-ancestry (British or non-British) UK Biobank target samples in most experiments, compared with *N*=320K in our main simulations, because the UK Biobank includes only 44K unrelated UK Biobank individuals of non-British European ancestry (we used *N*=293K unrelated British-ancestry UK Biobank target samples in a subset of experiments to more closely match our main simulations); (ii) we computed summary LD information using either *N*=400, *N*=4,000, or *N*=44K unrelated British-ancestry UK Biobank reference samples (either non-overlapping or overlapping with the target samples), or using *N*=3,567 reference samples from the UK10K cohort^35^ (compared with in-sample LD based on the target samples in the main simulations); (iii) we generated summary statistics using raw genotypes rather than summary LD information (as required when the target sample and the LD reference panel are not the same); (iv) we simulated 3 causal SNPs per locus that jointly explain 0.5% of trait variance, compared with 10 causal SNPs that jointly explain 0.05% of trait variance in our main simulations, to obtain sufficient power despite having a smaller sample size; and (v) in some experiments we used a subset of SNPs for generating causal SNPs or for fine-mapping analysis.

We now provide technical details of the differences between our simulations with mismatched reference LD and our main simulations:

- We simulated summary statistics using either 293,000 unrelated British-ancestry UK Biobank individuals, or 44,000 UK Biobank individuals of either British or non-British European ancestry, including related individuals in some experiments (using *N*=44,000 because this is the number of unrelated non-British European UK Biobank individuals). We determined ancestry based on self-reported ancestry, using UK Biobank data field 21000 (Ethnic background). We considered Irish-ancestry as a non-British European ancestry.
- We used LDStore^30^ v1.1 to compute summary LD information from UK10K sequencing data, and LDStore v2.0 to compute summary LD information from UK Biobank imputed genotypes.
- We generated summary statistics from imputed genotypes rather than summary LD information, using PLINK 2.0^75,76^.
- We performed each experiment four times, each time applying PolyFun + SuSiE while assuming either 1, 2, 3, or 10 causal SNPs per locus.

### Functionally informed fine-mapping of 49 complex traits in the UK Biobank

We applied SuSiE and PolyFun + SuSiE to fine-map 49 traits in the UK Biobank, including 31 traits analyzed in refs. ^38,39^, 9 blood cell traits analyzed in ref. ^12^, and 7 recently released metabolic traits (average *N*=318K; Supplementary Table 8), using the same data and the same parameter settings described in the Fine-mapping simulations section. We performed basic QC on each trait as described in our previous publications^38,39^. Specifically, we removed outliers outside the reasonable range for each quantitative trait, and quantile normalizing within sex strata after correcting for covariates for non-binary traits with non-normal distributions. We computed summary statistics with BOLT-LMM v2.3.3^39^ adjusting for sex, age and age squared, assessment center, genotyping platform, and the top 20 principal components (computed as described in ref. ^39^), and dilution factor for biochemical traits. As the non-infinitesimal version of BOLT-LMM does not estimate effect sizes, we computed z-scores for fine-mapping by taking the square root of the BOLT-LMM *χ*^2^ statistics and multiplying them by the sign of the effect estimate from the infinitesimal version of BOLT-LMM.

We partitioned each autosomal chromosome into 2,763 overlapping 3Mb-long loci with a 1Mb spacing between the start points of consecutive loci. We computed a PIP for each SNP based on the locus whose center was closest to the SNP (excluding SNPs >1Mb away from the closest center and loci wherein all SNPs had squared marginal effect sizes smaller than 0.00005). We excluded the MHC region (chr6 25.5M-33.5M) and two other long-range LD regions (chr8 8M-12M, chr11 46M-57M)^77^ from all analyses, following our observations that both methods tend to produce spurious results in these regions, finding many PIP=1 SNPs across many traits regardless of their BOLT-LMM p-values. We verified that other previously reported long-range LD regions^77^ do not harbor a disproportionate number of PIP>0.95 SNPs. We specified per-locus causal effect variances for SuSiE and PolyFun + SuSiE via our modified HESS approach. For all S-LDSC and fine-mapping analyses we specified a sample size corresponding to the BOLT-LMM effective sample size^39^ (given by the true sample size multiplied by the median ratio between *χ*^2^ statistics of BOLT-LMM and linear regression across SNPs with BOLT-LMM *χ*^2^>30).

All S-LDSC analyses used LD scores computed from in-sample summary LD information (based on imputed SNP dosages rather than sequenced genotypes as in previous publications^25–27^, assigning to each SNP the LD score computed in the locus in which it was most central) because they provide better coverage of low-frequency SNPs and are consistent with the fine-mapping analyses. We computed genetic correlations with LDSC, using the same summary statistics used for fine-mapping and restricting the analysis to common SNPs.

We selected a subset of 16 genetically uncorrelated traits by ranking all traits according to the number of PolyFun + SuSiE PIP>0.95 SNPs and greedily selecting top-ranked traits such that no selected trait has |*r_g_*|>0.2 with a previously selected trait, excluding traits having either (1) 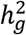 estimates <0.05 in either the PolyFun dataset (*N*=337K) or in the PolyLoc dataset (*N*=122K) (see 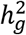 estimation description below); or (2) traits with an effective sample size <100K in the *N*=337K dataset (using 4/(1/#cases + 1/#controls) for binary traits).

We estimated 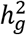 tagged by PIP>0.95 SNPs and by lead GWAS SNPs via a multivariate linear regression. We regressed all the covariates used in BOLT-LMM out of the phenotypes, performed multivariate linear regression on the residuals (using all PIP>0.95 SNPs as explanatory variables) and reported the adjusted *R*^2^ as the 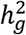 tagged by these SNPs. We verified that the results remained nearly identical regardless of whether we excluded related individuals (Supplementary Table 14). We estimated MAF>0.001 SNP-heritability for trait selection and for Figure 3b by running S-LDSC with all the baseline-LF annotations. We overrode the automatic removal of very large effect SNPs employed by S-LDSC for hair color, because this removal led to 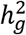 estimates that were smaller than the linear regression-based estimates, due to the large proportion of SNP-heritability originating from very large-effect SNPs.

We defined top annotations for Table 2, Figure 5 and Supplementary Tables 15-16 by first ranking all annotations according to their functional enrichment among PIP>0.95 SNPs (as in Figure 6; see below), and associating each SNP with its top ranked annotation, using meta-analyzed enrichment.

We selected a subset of genetically uncorrelated traits for each SNP (used in Figure 4, Table 2 and Supplementary Table 15), aiming to select traits from a diverse a set of groups as possible (anthropometric, lipids/metabolic, blood, cardiovascular/metabolic disease, other; Figure 4, Supplementary Table 8). To this aim, we iterated over trait groups cyclically. For each group containing ≥1 unselected traits with PIP>0.95 for the analyzed SNP, we selected the trait having the smallest average |*r_g_*| with unselected traits from other groups (if there remained any) or from all remaining traits (otherwise), selecting among all traits having |*r_g_*|<0.2 with previously selected traits, until no more eligible traits remained. We plotted the ideogram in Figure 4 with the PhenoGram^78^ software.

We computed enrichment of functional annotations among fine-mapped SNPs (Figure 6) as the ratio between the proportion of common SNPs with PIP above a given threshold having a specific annotation and the proportion of common SNPs having the annotation. We excluded continuous annotations and annotations constructed via windows around other annotations, and merged concordant annotations for common and low-frequency variants. We computed P-values using Fisher’s exact test (meta-analyzed across traits via Fisher’s method). We computed standard errors by (1) computing the standard error *s* of the log of the enrichment via the standard formula for the standard error of relative risk (exploiting the fact that enrichment and relative risk are both ratios of proportions); and (2) computing the standard error of the enrichment via 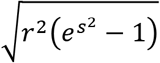 (i.e., the standard deviation of the exponent of a normal random variable), where *r* is the original enrichment estimate (meta-analyzed across traits using a fixed-effects meta-analysis). We excluded traits having <10 PIP>0.95 SNPs from the meta-analysis. The annotations shown in Figure 6 are non-synonymous, Conserved_LindbladToh (denoted Conserved), Human_Promoter_Villar_ExAC (denoted Promoter-ExAC), H3K4me3_Trynka (denoted H3K4me3), and Repressed_Hoffman (denoted Repressed).

To compare our fine-mapping results with those of refs.^7,12^, we restricted the comparison to SNPs that were not excluded from our fine-mapping procedure (SNPs having MAF≥0.001 in the UK Biobank *N*=337K dataset, INFO score≥0.6, distance <1Mb away from the closest locus center, and not residing in one of the excluded long-range LD regions). When the same SNP had multiple reported PIPs in ref. ^12^, we used the entry with the larger PIP. We caution that the comparison with ref. ^12^ is not a replication analysis because the datasets of ref. ^12^ and of PolyFun + SuSiE are correlated.

We selected traits for down-sampling analysis (analyzing *N*=107K individuals) as the set of traits having

(1) the largest number of 3Mb significant loci harboring a genome-wide significant SNP; (2) >10 PIP>0.95 SNPs in the SuSiE *N*=107K analysis; and (3) |*r_g_*|<0.2 with another selected trait.

### Polygenic localization

Polygenic localization aims to identify a minimal set of SNPs causally explaining a given proportion of common SNP heritability. To formally define polygenic localization, we first define expected common SNP heritability under a fixed effects model with no autocorrelation. We assume the linear model 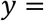 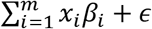, where *y* is a phenotype, *x*_*i*_ is a standardized common SNP with effect *β*_*i*_, *m* is the number of common SNPs and *ϵ* is a residual term. We define the expected common SNP heritability *Eh*^2^ as follows:

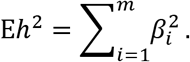

This definition stems from the fixed effects SNP-heritability 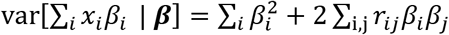, where we assumed standardized genotypes, *r*_*ij*_ is the LD between SNPs *i*, *j*, and the second term on the right hand size is zero in expectation, assuming no autocorrelation. We next define the expected common SNP heritability of a subset ***S*** of SNPs by:

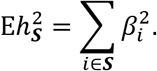

We define 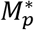, the smallest set of common SNPs causally explaining proportion *p* of expected common SNP heritability, as the cardinality of the smallest set ***S*** such that 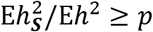. An equivalent definition is the smallest integer *k* such that

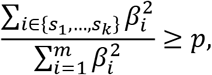

where *s*_*j*_ denotes a ranking of 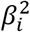 such that 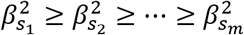. We analogously define *M*_*p*_ with respect to a given (possibly non-optimal) ranking of SNPs *s*′ as the smallest integer *k*′ such that 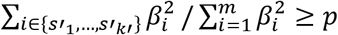. We note that 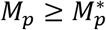 by construction.

Unfortunately, 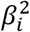 is unknown in practice. Polygenic localization therefore estimates an upper-bound of 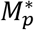 non-parametrically, by estimating *M*_*p*_ with respect to a ranking of SNPs based on PolyFun + SuSiE 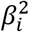 posterior mean estimates, using a random effects model. We first provide a brief conceptual description of PolyLoc and then describe it in detail below. Briefly, PolyLoc proceeds by (1) partitioning SNPs with similar 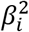 posterior mean estimates (using PolyFun + SuSiE estimates) into bins; (2) treating *β*_*i*_ as a zero-mean random variable and jointly estimating var[*β*_*i*_] in every bin using S-LDSC; and (3) finding the smallest integer *k* such that 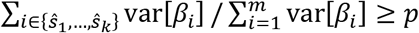, where 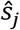 denotes the original ranking of 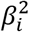 posterior mean estimates from PolyFun + SuSiE. The use of 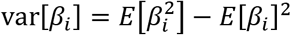 instead of 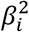 uses the assumption that *β*_*i*_ has zero mean in each bin. The partitioning into bins in step 1 induces a piecewise-linear approximation of the function 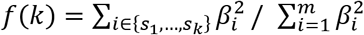. We use different datasets to estimate 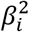 posterior means and to estimate var[*β*_*i*_] to prevent winner’s curse (which occurs when performing inference based on top ranked items using the same data used for ranking). Our approach is conservative by design due to using an imperfect ranking compared to the true ranking *s*_1_, …, *s*_*m*_. The degree of conservativeness is a function of fine-mapping power, and thus depends on factors affecting fine-mapping power such as sample size, levels of LD at causal SNPs, MAFs of causal SNPs, and trait polygenicity.

We now describe PolyLoc in detail. We used two sets of BOLT-LMM summary statistics based on different datasets: *N*=337,491 unrelated British-ancestry UK Biobank individuals, and *N*=121,768 European-ancestry UK Biobank individuals not included in the first set. PolyLoc proceeds as follows:

1. Apply median-based clustering of posterior mean estimates of 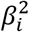 (i.e., the sum of the squared posterior mean and of the posterior variance of *β*_*i*_ reported by PolyFun + SuSiE, based on the PolyFun *N*=337K dataset) into 50 bins using the Ckmedian.1d.dp method^71^. Afterwards, include all SNPs excluded from fine-mapping (e.g. SNPs in the MHC region) in the last bin, and sub-partition the last bin into 10 equally-sized MAF bins to account for MAF-based genetic architecture^26^, yielding 59 bins (or 20 bins when also including non-common SNPs, yielding 69 bins). We used a larger number of bins than that used in step 3 of PolyFun because posterior mean estimates of 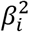 follow a heavy-tailed distribution, requiring many bins to avoid placing two SNPs having effect sizes of a different order of magnitude in the same bin.
2. Jointly estimate var[*β*_*i*_] of SNPs in each bin (defined as the SNP-heritability causally explained by the bin divided by the bin size) by creating an annotation for each bin and running a modified version of S-LDSC using the summary statistics from the PolyLoc *N*=122K dataset, with the following modifications: (a) use in-sample summary LD information (based on SNP dosages from imputed genotypes) to compute LD scores as in the PolyFun analyses; (b) apply non-negativity constraints to prevent negative var[*β*_*i*_] estimates; and (c) retain SNPs with *χ*^2^ > 80 in the analysis. Step (c) facilitates handling of bins that predominantly consist of very large effect SNPs.
3. Rank common SNPs according to their 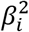 posterior mean estimates from PolyFun + SuSiE (if they do not reside in MAF bins) or according to a ranking of MAF bins (otherwise). To rank MAF bins, we applied standard S-LDSC to the PolyFun dataset (*N*=337K in our setting) with the same bin partitioning and ranked MAF bins according to their enrichment estimates (this is needed because 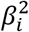 is not necessarily strongly correlated with MAF). Afterwards, compute *M*_**p**_ as the smallest number of top-ranked common SNPs, such that the sum of their var[*β*_*i*_] estimates from step 2 is equal to proportion *p* of the sum of all var[*β*_*i*_] estimates. Finally, compute standard errors of *M*_**p**_ via a 200-block-jackknife^79^, recomputing *M*_*i*_ separately using the estimates of each jackknife block.

Step 2 of PolyLoc requires a set of samples (*N*=122K in our analyses) different than that used in PolyFun + SuSiE (*N*=337K in our analyses) rather than a partitioning of chromosomes to avoid winner’s curse. Otherwise, PolyLoc would use the same summary statistics for both bin-partitioning and for estimating per-bin heritability (noting that this difficulty does not exist in PolyFun because PolyFun performs bin-partitions using only functional annotations, without requiring summary statistics from the target chromosome).

When computing standard errors of *M*_*p*_, we excluded jackknife blocks that yielded extremely noisy estimates (yielding an *M*_*p*_ estimate whose distance to the median of the estimates was >25x the interquartile range of all jackknife-block estimates; typically <2 blocks per trait). Such blocks likely result from the inclusion of very large-effect SNPs in step 2 of PolyLoc.

We included all MAF>0.001 SNPs in the set of S-LDSC regression SNPs (defined in refs. ^17,25,26^) regardless of whether we were interested in polygenic localization of common SNP-heritability or of MAF>0.001 SNP heritability, but did not include them in set of S-LDSC heritability SNPs (defined in refs. ^17,25,26^) except in analyses of MAF>0.001 SNP heritability.

In secondary analyses, we compared PolyLoc to an alternative method that performs polygenic localization based on prior estimates of per-SNP heritability from functional annotations, rather than posterior estimates. This alternative method uses per-SNP heritability estimates and SNP bins from step 4 of PolyFun, based only on the *N*=337K dataset (noting that it does not suffer from winner’s curse because PolyFun applies a partitioning into odd and even chromosomes).

PolyLoc will yield robust estimates of *M*_*p*_ if S-LDSC yields robust estimates of the SNP-heritability causally explained by each bin. Although S-LDSC has previously been shown to produce robust estimates^17,25–27^, we performed extensive simulations to confirm that PolyLoc produces robust estimates of *M*_*p*_. An exact simulation scheme would require first ranking all SNPs according to their PolyFun + SuSiE posterior per-SNP heritabilities, which is computationally prohibitive. To circumvent this computational challenge, we demonstrate that PolyLoc produces robust estimates of *M*_*p*_ with respect to several different SNP rankings. Specifically, we (1) generated causal effects for all SNPs on chromosome 1; (2) generated two independent corresponding sets of summary statistics for all SNPs on chromosome 1: A PolyFun dataset (with sampling noise based on *N*=320K individuals) and a PolyLoc dataset (with sampling noise based on *N*=122K individuals); (3) generated several different rankings of SNPs on chromosome 1 corresponding to different levels of statistical power (see below), using the PolyFun summary statistics (*N*=320K) from step 2; and (4) applied PolyLoc using the rankings from step 3, using the PolyLoc summary statistics from step 2 (*N*=122K). We generated causal effects and summary statistics in steps 1-2 as in the fine-mapping simulations (except for the restriction to chromosome 1), using LD matrices based on genotypes of 337K UK biobank individuals, such that 0.5% of the SNPs are causal and the SNPs jointly explain 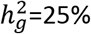. We ranked SNPs in step 3 according to approximate posterior per-SNP heritabilities given by 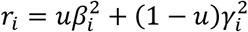, where *β*_*i*_ is the simulated causal effect of SNP *i*, [*γ*_1_, …, *γ*_*m*_] is a random permutation of [*β*_1_, …, *β*_*m*_], and *u* ∈ {0,0.25,0.5,0.75,1} represent power levels, such that *u* = 1 indicates maximal power and *u* = 0 indicates zero power. We partitioned SNPs into 10 bins in step 4 of each simulation (rather than 50 as in the real data analyses) because we only used chromosome 1 SNPs. (We verified that the results are relatively insensitive to the number of bins in secondary analyses). We generated 10 simulations for each evaluated value of *u*, and compared the estimated and true values of *M*_*p*_ in each simulation (using log scale because *M*_*p*_ spans different orders of magnitude for different levels of *p*).

PolyLoc yielded slightly conservative estimates of log_10_ *M*_*p*_ for *u*>0 and slightly anti-conservative estimates for *u*=0 (Supplementary Table 32). For *u*=0.75, representing a well-powered (but not perfectly-powered) study, the average bias of log_10_ *M*_50%_ was 0.007. We emphasize that we compare estimates of log_10_ *M*_*p*_ to their true values with respect to a given ranking (determined in step 3 of the simulation procedure) rather than the optimal ranking. We also compared log_10_ *M*_*p*_ estimates to 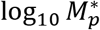 (reflecting the optimal ranking). As expected, the magnitude of the difference increased as *u* decreased. For example, the difference between the estimates value of log_10_ *M*_50%_ and 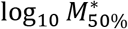 was 0.02 for *u*=1, 0.02 for *u*=0.75, 0.04 for *u*=0.5, 0.11 for *u*=0.25, and 2.9 for *u*=0 (Supplementary Table 32).

We performed 6 secondary analyses. First, we repeated the analysis using a version of S-LDSC that does not constrain per-SNP heritabilities to be non-negative. The results were similar (Supplementary Table 32), but interpretation was more challenging because some per-SNP heritability estimates became negative. Second, we varied the PolyFun sample size in the range 107K - 1 million and obtained qualitatively similar results (Supplementary Table 32). Third, we varied the SNP heritability in the range 12.5%-50% and obtained qualitatively similar results (Supplementary Table 32). Fourth, we varied the genome-wide proportion of causal SNPs in the range 0.1%-1% and obtained qualitatively similar results (Supplementary Table 32). Fifth, we varied the number of bins used for partitioning SNPs in the range 5-20 and obtained qualitatively similar results (Supplementary Table 32). Finally, we evaluated the results with respect to a ranking of SNPs based only on the magnitude of their summary statistics. We obtained highly conservative results with respect to both the true value of *M*_*p*_ and to 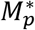 in all cases (Supplementary Table 33), demonstrating that accurate polygenic localization estimates require performing genome-wide fine-mapping rather than simply ranking SNPs based on their summary statistics.

## Acknowledgements

We thank Bogdan Pasaniuc, Gleb Kichaev, Matthew Stephens, Gao Wang, Masahiro Kanai, Brian M. Schilder and Towfique Raj for helpful discussions. This research was conducted using the UK Biobank Resource under Application #16549 and was funded by NIH grants U01 HG009379, R37 MH107649, R01 MH101244 and R01 HG006399 HG006399, and by the Academy of Finland grants 288509 and 312076. HKF is supported by Eric and Wendy Schmidt. Computational analyses were performed on the O2 High-Performance Compute Cluster at Harvard Medical School.

## URLs

Software implementing PolyFun and PolyLoc: https://www.hsph.harvard.edu/alkes-price/software

Baseline-LF v2.2.UKB annotations and LD-scores for UK Biobank SNPs: https://data.broadinstitute.org/alkesgroup/LDSCORE/baselineLF_v2.2.UKB.tar.gz

Summary LD information of *N*=337K British-ancestry UK Biobank individuals for 2,673 overlapping 3Mb loci: https://data.broadinstitute.org/alkesgroup/UKBB_LD/

Fine-mapping results for all analyzed SNPs: https://data.broadinstitute.org/alkesgroup/polyfun_results/

SuSiE: https://github.com/stephenslab/susieR

FINEMAP: http://www.christianbenner.com/#

UK Biobank Resource: http://www.ukbiobank.ac.uk/

## Code and data availability

PolyFun and PolyLoc are open-source software packages freely available at https://github.com/omerwe/polyfun. Access to the UK Biobank resource is available via application (http://www.ukbiobank.ac.uk/). PolyFun fine-mapping results generated in this study are currently available for public download at http://data.broadinstitute.org/alkesgroup/polyfun_results. Summary LD information generated in this study are available for public download at https://data.broadinstitute.org/alkesgroup/UKBB_LD

