## Supplementary Material for "Functionally-informed fine-mapping and polygenic localization of complex trait heritability"

### Supplementary Information for: Functionally-informed fine-mapping and polygenic localization of complex trait heritability

#### Supplementary Table Captions

**Table S1: Description of baseline-LF model annotations**. For each annotation we report #SNPs (unless it is a continuous-valued annotation), #common SNPs, whether it is binary or continuous-valued, and a literature reference.

**Table S2: Description of 10 3Mb loci used in simulations**. We report the start and end position of each locus (using hg19 coordinates), as well as the number of SNPs in that locus.

**Table S3: Functional enrichments used to generate simulated data**. For each annotation, we report the coefficient $\tau$ used in the generative model, as computed via a meta-analysis across 16 real traits.

**Table S4: Numerical results of fine-mapping simulations**. Each row reports simulation results for a unique combination of a method, method settings, simulation settings and a PIP cutoff. The columns are (**A**) Method: Method name; (**B**) Functional: the type of functional data used (None: non-functionally informed analysis; PolyFun: PolyFun; default: default settings for CAVIARBF and fastPAINTOR; S-LDSC priors: priors from standard S-LDSC; S-LDSC+L2 priors: Priors from L2-regularized S-LDSC; cheat: fine-mapping with true prior causal probabilities; PolyFun (no L2): PolyFun that does not apply L2-regularization); (**C**) PIP cutoff: The PIP cutoff evaluated; (**D**) N: simulated sample size; (**E**) h2: The SNP-heritability causally explained by SNPs at the target locus; (**F**) c: number of simulated causal SNPs at the target locus; (**G**) functional architecture: the function that relates functional annotations to prior causal probabilities (additive, multiplicative, or sub-additive); (**H**) q: the ratio between the highest and lowest prior causal probability; (**I**) Max #causal SNPs: The maximum number of causal SNPs allowed (or the exact number for SuSiE and PolyFun + SuSiE); (**J**) Genome-wide h2: the SNP-heritability causally explained by all genome-wide SNPs; (**K**) Genome-wide sparsity: The proportion of genome-wide non-causal SNPs; (**L**) Max #annotations: The maximum number of annotations allowed (only relevant for fastPAINTOR); (**M**) variance estimator: determines whether the (method-specific) default estimator or the HESS-based estimator of causal effect size variance was used for fine-mapping; (**N**) FDR: false discovery rate under the evaluated PIP cutoff; (**O**) FDR (s.e.): the standard error of the FDR (evaluated via jackknife); (**P**) power: empirical power under the evaluated PIP cutoff; (**Q**) power (s.e.): the standard error of the power (evaluated via jackknife); (**R**) #experiments: The number of experiments performed for this combination of settings; (**S**) Avg. #fine-mapped SNPs: The average number of SNPs with PIP greater than the cutoff; (**T**) FDR threshold: the exact FDR threshold, given by one minus the average PIP among SNPs with PIP greater than the cutoff; (**U**) Time (avg): The average time required to perform fine-mapping (minutes); (**V**) Time (s.d.): The standard deviation of time required to perform fine-mapping (minutes); (**W**) 95% CS size: The size of the union of all 95% credible sets, averaged across simulations (only relevant for FINEMAP and SuSiE-based methods); (**X**) %Causal SNPs in 95% CS: The proportion of causal SNPs found in the union of all 95% credible sets (averaged across simulations).

**Table S5: Sensitivity of PolyFun method to perturbing steps 1-5.** For each of step 1 (estimating functional enrichments), step 2 (estimating per-SNP heritabilities on odd/even chromosomes), step 3 (estimating per-SNP heritabilities on odd/even chromosomes), step 4 (re-estimating per-SNP heritabilities within each bin excluding the target chromosome) and step 5 (specifying prior causal probabilities in proportion of the re-estimated per-SNP heritabilities), we report the perturbation that we implemented; the impact on false positives (FDR), if statistically significant; and the impact on power, if statistically significant. *denotes significant change in results only in simulations with smaller sample size (N=10K).

**Table S6: Numerical results of simulations that assess the impact of each of PolyFun steps (1)-(5) on fine-mapping performance.** The table is similar to Table S4 with several new columns: (**1**) Table: table alphabetic reference; **(2)** %flipped annot. coeff.: the proportion of annotation coefficients whose sign was flipped in step 1; **(3)** Odd/even split – whether PolyFun steps 1 and 2 handled even and odd chromosomes separately; **(4)** #bins – the number of SNP bins into which PolyFun assigns SNPs with similar predicted per-SNP heritabilities in step 3; **(5)** Exclude target chrom.: whether the target chromosome was excluded from the re-estimation of per-SNP heritabilities in step 4; **(6)** %permuted priors – the proportion of SNPs in the target locus whose prior causal probability was randomly swapped with that of another SNP.

**Table S7: Numerical results of simulations of mismatch between the GWAS sample and the LD reference panel.** The table is similar to Table S4, with constant columns omitted (e.g. genome-wide $h^2$, which is equal to the default simulation value); and several new columns: (**1**) Table: table alphabetic reference; (**2**) GWAS sample: The population from which GWAS individuals who are not in the LD reference panel were selected (for overlapping individuals, the relevant column is "LD sample") (UK: unrelated British-ancestry UK Biobank individuals; EUR: unrelated non-British European-ancestry UK Biobank individuals; UK-REL: British-ancestry UK Biobank individuals without excluding pairs of related individuals; EUR-REL: Non-British European-ancestry UK Biobank individuals without excluding pairs of related individuals); (**3**) LD sample: The population from which the LD reference panel was sampled (UK: unrelated British-ancestry UK Biobank individuals; UK10K: The UK10K cohort); (**4**) N GWAS: The GWAS sample size; (**5**) N LD: The LD reference panel sample size (0 indicates that we used an LD matrix that is equal to the identity matrix); (**6**) N overlap: The number of individuals in the GWAS sample who are also included in the LD reference panel (noting that they can be from a different population from the population reported in the "GWAS sample" column); (**7**) Generative SNPs: the set of SNPs from which causal SNPs were selected; (**8**) SNPs considered: The set of SNPS included in fine-mapping.

**Table S8: Description of 49 UK Biobank traits analyzed.** For each trait we report its group (used in Figure 4); whether it is binary or continuous-valued; sample size analyzed by PolyFun (max N=337K unrelated British-ancestry samples) and PolyLoc (max N=122K non-overlapping samples); the number of cases in the PolyFun and PolyLoc sample sets (for binary traits); and the heritability causally explained by MAF>0.001 SNPs (using observed-scale heritability for binary traits), and its standard error, in the PolyFun dataset (max N=337K) and in the PolyLoc dataset (max N=122K).

**Table S9: Comparison of PolyFun + SuSiE vs. SuSiE fine-mapping results for UK Biobank traits**. We report the number of fine-mapped SNPs found by PolyFun + SuSiE and by SuSiE for various trait and PIP cutoffs. For every trait and one of three evaluated PIP cutoffs (0.05 0.5, 0.95) we report the number of SNPs with PIP greater than this cutoff identified by PolyFun + SuSiE, SuSiE, and by both (i.e. SNPs at the intersection of these two SNP sets).

**Table S10: List of all fine-mapped SNPs for UK Biobank traits**. We report the list of all SNPs with PIP>0.95 in either PolyFun + SuSiE or SuSiE. For every SNP we report its rsid, position (using hg19 coordinates), major (A1) and minor (A2) alleles, PIP, posterior per-SNP $h^2$ (i.e., the sum of the squared posterior mean and posterior variance of its causal effect size), the posterior mean and standard deviation (sd) of its cal effect size, the index of one of the 95% credible sets to which it belongs under each of the two methods, its MAF in the entire UK Biobank and in the N=337K dataset we analyzed, its distance to the nearest lead GWAS SNP, the center of the locus in which it was analyzed, its BOLT-LMM p-value, its BOLT-LMM marginal effect size estimate (BETA), its prior causal probability, and whether it is coding and non-synonymous.

**Table S11: Genetic correlations between UK Biobank traits.** We report a matrix of $r_g$ estimates, computed using N=337K UK Biobank individuals (see Methods).

**Table S12: List of 16 genetically uncorrelated UK Biobank traits.** We report the 16 genetically uncorrelated traits selected for many of our primary analyses. All the traits have pairwise $|r_g|$<0.2, $h_g^2$>0.05 in the two different subsets of the UK Biobank, and an effective sample size >100K in the N=337K data set (see Methods).

**Table S13: Distribution of PIP>0.95 SNPs across GWAS loci for UK Biobank traits**. For each lead GWAS SNP, we report the number of PIP>0.95 SNPs within distance of at most 0.5Mb from the lead SNP.

**Table S14: Numerical values of heritability tagged by PIP>0.95, heritability tagged by fine-mapped SNPs, and heritability causally explained by MAF>0.001 SNPs for UK Biobank traits.** For each of the 16 genetically uncorrelated traits we report the heritability causally explained by MAF>0.001 SNPs (as estimated via S-LDSC), the heritability tagged by PIP>0.95 SNPs (estimated via the adjusted $R^2$ of a multivariate linear regression) and the heritability tagged by lead GWAS SNPs (estimated via the adjusted $R^2$ of a multivariate linear regression), using observed-scale heritability for binary traits. All analyses were performed using a set of (max N=122K) individuals not analyzed by PolyFun + SuSiE, to prevent winner's curse. We also report the corresponding estimates obtained via a multivariate linear regression using a subset of N=45K unrelated individuals, which were nearly identical to the N=122K results.

**Table S15: List of all pleiotropic fine-mapped SNPs (PIP>0.95) identified by PolyFun + SuSiE for UK Biobank traits.** The table is analogous to Table 2 but includes results for all 223 PIP>0.95 pleiotropic SNPs. The nearest gene column was specified via (1) the GWAS catalog "mapped gene" entry if the SNP appeared in the GWAS catalog; (2) the dbSNP gene entry for the SNP, if such an entry exists; or (3) a manual inspection in the UCSC genome browser. The nearest gene column for intergenic SNPs sometimes includes two genes separated by a dash, representing the two closest flanking genes.

**Table S16: Examples of the advantages of functionally-informed fine-mapping for UK Biobank traits.** For all 121 loci where PolyFun + SuSiE identified a PIP>0.95 SNP and SuSiE did not identify a single PIP>0.5 SNP within 500kb of that SNP, we report the prior causal probability and annotations of the fine-mapped SNP and of the top SuSiE SNP in this locus.

**Table S17: Numerical values of functional enrichment of SuSiE fine-mapped common SNPs for UK Biobank traits**. For each main functional annotation (Methods), PIP range, and trait, we report enrichment, its standard error, and its p-value. We report meta-analysis results in separate rows with the trait name "Meta-analysis".

**Table S18: Summary of comparison between our results and those of Ulirsch *et al.* 2019 Nat Genet**. The table reports the number of SNPs fine-mapped (PIP>0.95) by Ulirsch *et al.* for 9 blood cell traits, the number of SNPs fine-mapped (PIP>0.95) by one of our evaluated methods/datasets (a different method/dataset in every row) and the size of the intersection of the two sets.

**Table S19: Detailed comparison between our results and those of Ulirsch *et al.* 2019 Nat Genet**. The table includes the identities and PIPs of each SNP having PIP>0.95 in at least one of the evaluated methods.

**Table S20: Detailed comparison between our results and those of Ulirsch *et al.* 2019 Nat Genet for four SNPs that were functionally validated via luciferase reporter assays**. The table includes a subset of Table S19 for only the four SNPs of interest.

**Table S21: Summary of comparison between our results and those of Farh et al. 2015 Nature**. The table is analogous to Table S18 but compares our results to Farh *et al.* rather than Ulirsch *et al.*

**Table S22: Detailed comparison between our results and those of Farh et al. 2015 Nature.** The table is analogous to Table S19 but compares our results to Farh *et al.* rather than Ulirsch *et al.*

**Table S23: Comparison of PolyFun + SuSiE vs. SuSiE results for UK Biobank traits when down-sampling the data to N=107K individuals from the UK Biobank interim release**. For each of five selected traits (Methods) and all combinations of pairs of analyses (SuSiE or PolyFun + SuSiE, using either N=337K or N=107K subsets of the UK Biobank), we report the number PIP>0.95 SNPs identified by each method, and the number of PIP>0.95 SNPs identified by both methods.

**Table S24: Comparison of PolyFun + SuSiE vs. SuSiE 95% credible set sizes across all locus-trait pairs for UK Biobank traits**. We report the sizes of 95% credible sets for SuSiE and PolyFun + SuSiE for each locus-trait pair analyzed.

**Table S25: List of pairs of coding and non-coding fine-mapped SNPs within 1Mb of each other**. We report all pairs of fine-mapped coding and non-coding SNPs within 1Mb of each other, which could aid in linking regulatory variants to genes.

**Table S26: Numerical values of functional enrichment of PolyFun + SuSiE fine-mapped common SNPs for UK Biobank traits**. The table is analogous to Table S17 but uses PolyFun + SuSiE instead of SuSiE to compute PIPs.

**Table S27: Numerical values of functional enrichment of SuSiE fine-mapped MAF>0.001 SNPs for UK Biobank traits**. The table is analogous to Table S17 but uses MAF>0.001 SNPs instead of only common (MAF>0.05) SNPs.

**Table S28: Numerical values of functional enrichment of SuSiE fine-mapped low-frequency and rare SNPs for UK Biobank traits**. The table is analogous to Table S17 but uses 0.05>MAF>0.001 SNPs instead of common (MAF>0.05) SNPs.

**Table S29: Numerical polygenic localization results for 49 UK Biobank traits.** For each trait we report $M_p$ estimates (and their standard errors) at multiple values of $p$.

**Table S30: Comparison between polygenic localization and $M_e$ estimates for 14 UK Biobank traits**. We report $M_e$ and $M_{50\%}$ estimates for 13 genetically uncorrelated traits for which there exist published $M_e$ estimates (from O'Connor et al. 2019).

**Table S31: Polygenic localization results for secondary analyses of UK Biobank traits**. The table reports the results of all the secondary analyses described in the polygenic localization section of the paper. The analysis type is specified in the "Analysis" column.

**Table S32: Results of polygenic localization simulations**. Each row contains results for a specific combination of settings: u – simulated power (u=1 means perfect power and u=0 means no power; see Methods); h2 – simulated SNP-heritability; p – simulated proportion of causal SNPs; n – simulated sample size for the PolyFun dataset; num_bins – the number of bins used in the simulation; logM*p_mean - the number of SNPs causally explaining proportion *p* of SNP-heritability under the optimal ranking, averaged across 10 simulations; logM*p_sd – the standard deviation of logM*p across 10 simulations; logMp_mean – the average estimate of logMp across 10 simulations; logMp_sd – the standard deviation of estimates of logMp under 10 simulations; logMp_diff – the average difference between the estimates and true values of logMp under 10 simulations (positive values indicate conservative estimates); logMp_sd – the standard deviation of the difference between the estimates and true values of logMp across 10 simulations. All the logarithm in this table are log10 based. We note that logMp_diff is evaluated with respect to the true value of logMp rather than logM*p (see Methods).

**Table S33: Results of polygenic localization simulations using ranking based on magnitude of summary statistics.** The table is analogous to Table S32 but uses a ranking of SNPs based solely on the magnitude of their summary statistics.

### Supplementary Figures

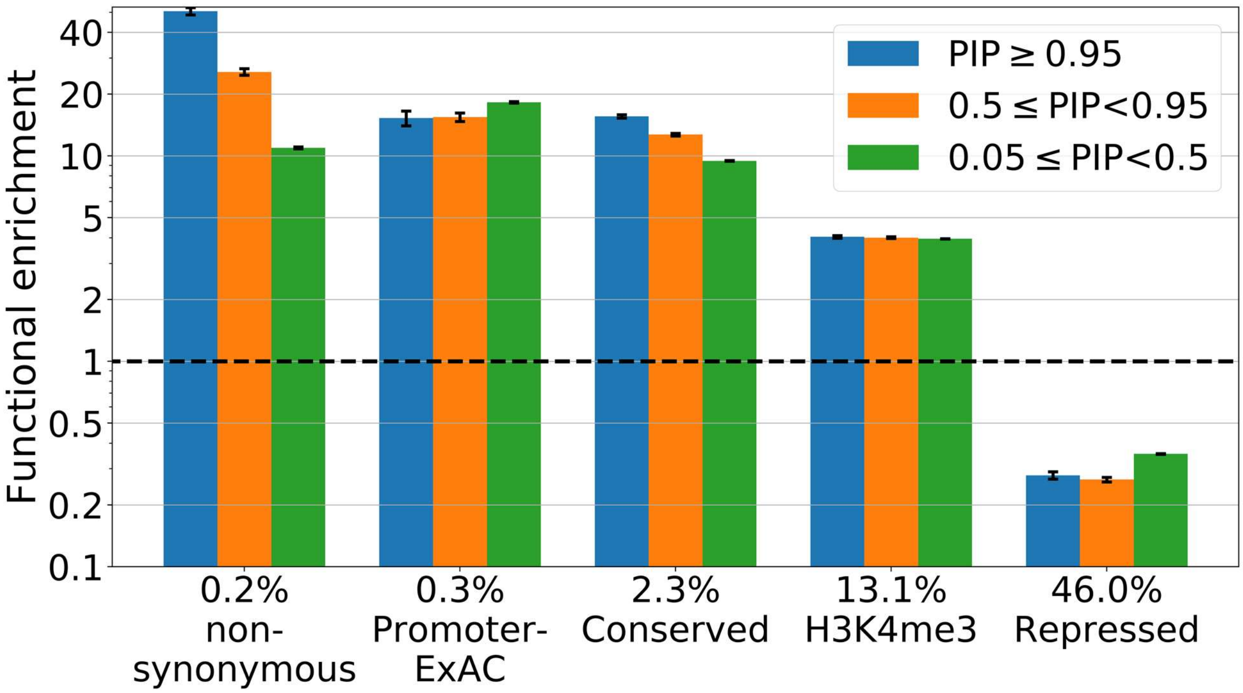


**Figure S1: Functional enrichment of PolyFun + SuSiE fine-mapped common SNPs for UK Biobank traits.** The figure is analogous to Figure 6 but uses PIPs computed by PolyFun + SuSiE instead of SuSiE.

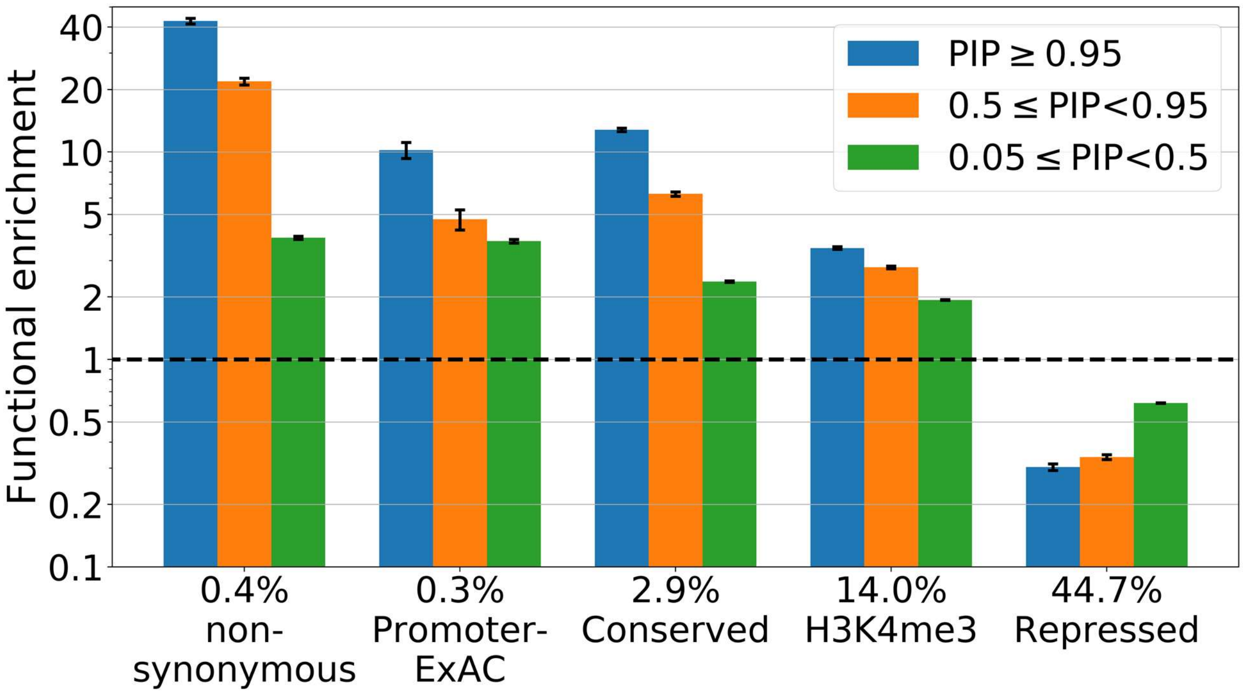


**Figure S2: Functional enrichment of SuSiE fine-mapped MAF>0.001 SNPs for UK Biobank traits.** The figure is analogous to Figure 6 but uses MAF>0.001 SNPs instead of common (MAF>0.05) SNPs.

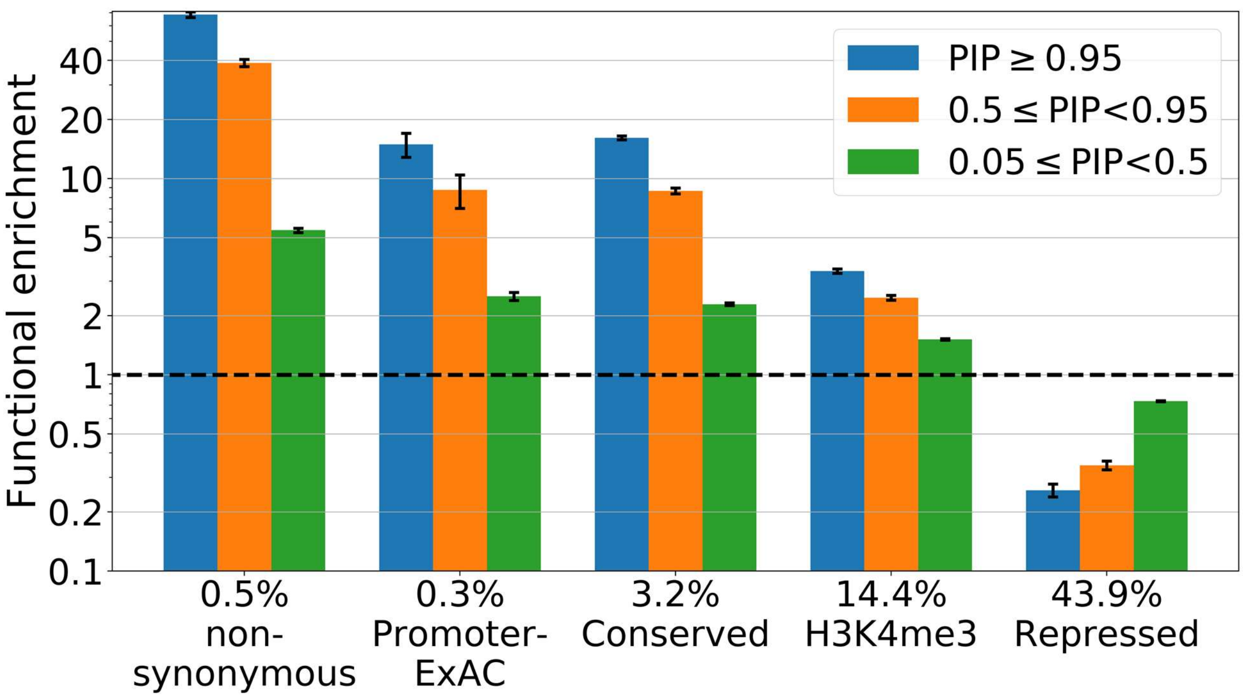


**Figure S3: Functional enrichment of SuSiE fine-mapped low-frequency and rare SNPs for UK Biobank traits.** The figure is analogous to Figure 6 but uses only low-frequency and rare SNPs (0.05>MAF>0.001) instead of common (MAF>0.05) SNPs.

### Supplementary Note

#### PolyFun + SuSiE does not make use of summary LD information when restricting to a single causal SNP per locus

We prove here that when applying PolyFun + SuSiE assuming that there $L$ causal SNPs per locus, the model only requires inverting $L \times L$ LD matrices of subsets of $L$ SNPs. Consequently, the model does not require an LD matrix under $L = 1$, because the LD of an SNP with itself is 1.0 by construction. Our derivation also supports the empirical result that PolyFun + SuSiE is relatively robust to LD misspecification under $L = 2$, presumably because the inverse of a $2 \times 2$ misspecified LD matrices is robust to moderate LD misspecification.

We first establish notations. Denote $\boldsymbol{z}$ as a $m \times 1$ vector of z-scores of $m$ SNPs, $\boldsymbol{R}$ as an $m \times m$ LD matrix, and $\beta_i$ as the causal effect of SNP $i$. Given a set of causal SNPs $\boldsymbol{C}$, define $\mathrm{P}(\boldsymbol{C})$ as the prior probability of the SNPs in set $\boldsymbol{C}$ to be causal: $\mathrm{P}(\boldsymbol{C}) = \prod_{i \in \boldsymbol{C}} \mathrm{P}(\beta_i \neq 0)$. Finally, define $\mathrm{BF}(\boldsymbol{C})$ as the Bayes factor of set $\boldsymbol{C}$, defined as $\mathrm{BF}(\boldsymbol{C}) = \mathrm{P}(\boldsymbol{z} \mid \boldsymbol{C}, \boldsymbol{R}) / \mathrm{P}(\boldsymbol{z} \mid \boldsymbol{R}; \boldsymbol{0})$, where $\mathrm{P}(z \mid \boldsymbol{R}; \boldsymbol{0})$ is the density of $\boldsymbol{z}$ assuming that there are no causal SNPs in the locus.

When applying PolyFun + SuSiE assuming $L$ causal SNPs per locus, the PIP of SNP $i$ is given by:

$$
\begin{aligned}
\mathrm{PIP_i} &= \mathrm{P}(\beta_i \neq 0 \mid \boldsymbol{z}, \boldsymbol{R}; L) \\
&= \mathrm{P}(\beta_i \neq 0\,) \frac{\mathrm{P}(\boldsymbol{z} \mid \beta_i \neq 0, \boldsymbol{R}; L)}{\mathrm{P}(\boldsymbol{z} \mid \boldsymbol{R}, L)} \\
&= \frac{\sum_{\mathbf{C}, |\boldsymbol{C}| = L, i \in \boldsymbol{C}} \mathrm{P}(\boldsymbol{C}) \mathrm{P}(\boldsymbol{z} \mid \boldsymbol{C}, \boldsymbol{R})}{\sum_{\mathbf{C}', |\boldsymbol{C}'| = L} \mathrm{P}(\boldsymbol{C}') \mathrm{P}(\boldsymbol{z} \mid \boldsymbol{C}', \boldsymbol{R})} \\
&= \frac{\sum_{\mathbf{C}, |\boldsymbol{C}| = L, i \in \boldsymbol{C}} \mathrm{P}(\boldsymbol{C}) \mathrm{BF}(\boldsymbol{C}) \mathrm{P}(z \mid \boldsymbol{R}; \boldsymbol{0})}{\sum_{\mathbf{C}', |\boldsymbol{C}'| = L} \mathrm{P}(\boldsymbol{C}') \mathrm{BF}(\boldsymbol{C}') \mathrm{P}(z \mid \boldsymbol{R}; \boldsymbol{0})} \\
&= \frac{\sum_{\mathbf{C}, |\boldsymbol{C}| = L, i \in \boldsymbol{C}} \mathrm{P}(\boldsymbol{C}) \mathrm{BF}(\boldsymbol{C})}{\sum_{\mathbf{C}', |\boldsymbol{C}'| = L} \mathrm{P}(\boldsymbol{C}') \mathrm{BF}(\boldsymbol{C}')}.
\end{aligned}
$$

We complete the proof by noting that $\mathrm{BF}(\boldsymbol{C})$ depends only on the $|\boldsymbol{C}| \times |\boldsymbol{C}|$ submatrix of $\boldsymbol{R}$ corresponding to SNPs in the set $\boldsymbol{C}$, as shown in previous work[1,2].

We conclude that $\mathrm{PIP_i}$ depends only on $L \times L$ submatrices of the matrix $\boldsymbol{R}$.
